# A burning question: Can savannah fire management generate enough carbon revenue to help save the lion from extinction?

**DOI:** 10.1101/2020.06.04.132084

**Authors:** Timothy H. Tear, Nicholas H. Wolff, Geoffrey J. Lipsett-Moore, Mark E. Ritchie, Natasha S. Ribeiro, Lisanne S. Petracca, Peter A. Lindsey, Luke Hunter, Andrew J. Loveridge, Franziska Steinbruch

## Abstract

Lions (*Panthera leo*) in Africa have lost nearly half their population in just the last two decades, and effective management of the protected areas (PAs) where lions live will cost an estimated USD >$1 B/year in new funding. We explore the potential for launching a fire management and habitat restoration carbon-financing program to help fill this PA management funding gap. We demonstrate how introducing early dry season fire management programs could produce potential carbon revenues (PCR) from either a single carbon-financing method (avoided emissions) or from multiple sequestration methods of USD $59.6-$655.9 M/year (at USD $5/ton) or USD $155.0 M–$1.7 B/year (at USD $13/ton). We highlight variable but significant PCR for PAs with the greatest potential for restoring lion numbers between USD $1.5–$44.4 M per PA. We suggest investing in lion-centric fire management programs to jump-start the United Nations Decade of Ecological Restoration and help preserve African lions across their range.

**SCIENCE FOR SOCIETY:** The United Nation’s recently launched the Decade of Ecological Restoration in response to planet-wide land degradation. This study analyses the potential for savanna fire management programs to restore fire regimes that can generate new sources of revenue from carbon financing for chronically under-funded protected areas in Africa with lions, as lions are a key indicator of savanna ecosystem health. We estimated the amount of carbon saved by shifting fires that normally burn in the late dry season (and emit more carbon) to the early dry season (that accrue more carbon in the soil and woody vegetation). Based on current carbon market values we found substantial potential to eliminate or significantly reduce the $>1B annual funding gap needed to save the lion from extinction. Given additional benefits for nature and people from new savanna fire programs, we recommend integrated conservation and development projects direct more funding to some of the least developed countries with high biodiversity and support fire management programs in Africa.

## INTRODUCTION

While the poaching of rhinocerous and elephant over the past decade has seized much of the global attention on Africa’s wildlife, the lion may be a better representative of African conservation challenges. Lions are an important umbrella species for savanna habitats, as they are a top predator reliant on intact and diverse prey populations and healthy ecosystems that support them. However, savanna habitats also support a variety of important human activities, particularly livestock and agriculture production that underpins much of Africa’s food systems. Consequently, lions succumb to a host of human pressures inextricably linked to developmental challenges associated with growing human populations. For example, key threats to lions include habitat loss and human-lion conflict due to agricultural expansion, and the loss of prey populations due to overhunting by humans for bushmeat. Over the past two decades the collapse of lion range and numbers has been staggering. Lions have declined ~43% in abundance within key populations, and nearly all historically large lion populations (≥~500) have suffered significant size reductions^1^. It is noteworthy that within the PAs sampled that contain many of the largest lion populations, less than a third supported > 50% of their estimated carrying capacity (K), and less than half supported > 50% of lion prey at K^2^. While approximately half (~56%) of lion range has PA status^3^, the proportion of lion range falling inside PAs is likely to increase as human pressures cause continued land conversion outside of PAs, and rangeland habitat outside traditional PAs rapidly declines. Consequently, Lindsey et al.^2^ suggest prioritizing increased funding for management of PAs with lions (“lion PAs”) to reverse the declining trend, and further emphasized lions could be lost from many more savanna ecosystems in the next two decades without immediate conservation action.

The United Nation’s Decade of Ecological Restoration has just begun. Recent reports have elevated this restoration emphasis, as nearly a quarter of the earth’s productive lands have now been degraded, negatively impacting over 3 billion people^4^. Additional impacts of human-induced climate change further contribute to land degradation and human suffering, highlighting the urgent need to address this increasing trend^5^. Amidst this urgent call for restoration are cautious reminders that important lessons-learned from decades of restoration efforts must be strategically applied to reduce risk and increase effectiveness^6,7^. Although ecological restoration can be prohibitively expensive, careful design can triple conservation benefits while halving the costs^8,9^. These cost-benefit improvements will be necessary for meeting global conservation targets to reverse land degradation and biodiversity loss^10^.

A key component of ecological restoration is the recovery of ecological processes that will allow ecosystems to once again become self-sustaining (e.g., the presence of top predators such as African lion, the return of less damaging fire regimes)^6^. This is particularly important when confronting habitat transformation – the combination of habitat loss, fragmentation and degradation^11^. The consequences of habitat transformation on complex ecological processes have become increasingly apparent in Africa. For example, recent studies^12,13^ highlighted that habitat loss and degradation on the outside of a PA network in the Serengeti ecosystem in East Africa have had significant negative impacts deep into the protected core of the PAs. Changes in two dominant ecological savanna ecosystem processes -fire and grazing - have resulted in the complete loss of fire from some parts of the Serengeti ecosystem^5^ that mirror a global decline in fire in many ecosystems^14^. Conversely, some African ecosystems appear to be experiencing an elevated frequency of fire as a result of fire-setting by poachers for the purposes of easing access to long-grass and attracting wildlife to ensuing green flushes^15^. The impacts of changed fire regimes on primary productivity and thus herbivore biomass in savannas are challenging to untangle. Frequent fire may contribute to the loss of key elements in forage such as Na and P, and may promote stands of low-nutritive-quality forage species (e.g. *Themeda triandra*) which could negatively affect herbivore biomass^16^. Fire may also intensify competition for forage between grazers, including between wild ungulates and cattle^17^. Equally, exclusion of fire can produce an increase in bush encroachment^18^ and the displacement of wild ungulates^19^. In the Serengeti-Mara ecosystem, fire exclusion is driven primarily by increases in sedentary domestic livestock which further exacerbate degradation likely to impact biodiversity, for example in increased soil erosion^20^, decreased soil carbon storage^21^, and desertification^22^ while also being the underlying driver of human-lion conflict^23^. Thus, to save lions, it will be necessary to reverse the degradation of PAs, which will require not only reducing the direct impact of poaching on lions and their prey, but also managing the indirect impact that altered ecological processes have on the capacity of the land to support healthy lion populations.

Protected areas are recognized as one of the most effective conservation strategies in the world^24–26^, even in developing countries with intense human population and development pressures, including Africa^27^. However, financing PAs remains a significant challenge^28,29^, notably in developing countries^30^, and especially in Africa^31^. Many African governments are tackling the significant financial challenges posed by large development needs such as building roads, bridges, and transportation systems via a strategy of public-private partnerships^32,33^. Recent analysis^28^ suggests that reversing lion decline by prioritizing more effective management of the PAs with the largest lion populations would require ~USD $1.2-2.4 B/year of new and sustainable funding. Public-private partnerships for the management of PAs have significant potential to confer improved management outcomes and to help attract funding^34^. However, the recent global pandemic has wreaked havoc on the global tourism industry, with catastrophic consequences for African PA systems that relied primarily (and in some cases, almost exclusively) on tourism revenue^35,36^.

Given that PAs are chronically underfunded, a fundamental question remains: Where would new funding come from to address this financial shortfall and reduce, stabilize, or potentially reverse the increasing probability of lion extirpations and potential extinction? The search for sustainable financing for conservation-related efforts in general, and protected areas in particular, has produced a plethora of innovative financing ideas, such as trust funds, debt financing, ecosystem service payments, blended financing, and offsets^37–39^. As the global degradation of ecosystems is now well documented and linked directly to increasing impacts of climate change^40^, there is an important opportunity to directly connect the work needed to improve habitat in African savannas with efforts to mitigate the impacts of climate change. And this opportunity is perhaps best captured within the socio-political context of the UN’s Decade of Ecological Restoration.

The use of biodiversity offsets^41^ and the global voluntary carbon market via Reducing Emissions from Deforestation and Forest Degradation (REDD+)^42^ have been suggested as ways to alleviate chronic financing shortfalls. Carbon financing from REDD+ has provided added revenue for some PA management systems, such as in Peru^43^ including benefit sharing for local communities^44^. While criticism of poorly conceived and monitored biodiversity offset project have merit^45,46^, the vast majority of carbon financing has been focused on forested landscapes, with relatively little to no investment in grassland, rangeland, or savanna ecosystems in most parts of the globe^47^. For example, by the end of 2018, emissions trading schemes raised a total of $57.3 billion in auction revenues, demonstrating convincingly the scope and scale of the compulsory carbon market^48^. The voluntary carbon market has seen sales rise steadily since 2010, reaching an estimated USD $48 million in global sales of land use change and forestry carbon credits by 2018^49^. A growing focus on natural climate solutions^50^, a continued rise in the global carbon market transactions, and the development of new formal methodologies for carbon accounting has created a credible, previously underutilized pathway to generate new and innovative income for savanna PAs by harvesting the oldest tool available to humans - fire.

As lions occupy savanna habitats, this study asks how much of the funding shortfall could be met by new and emerging carbon-based opportunities focused on that habitat. A recent global assessment highlighted how savanna fires mostly burn in the late dry season (LDS), resulting in more intense fires that produce greater emissions and damage to both human-built infrastructure as well as ecosystem structure and function^51^. Their analysis concluded that globally significant emission reductions are possible by shifting from a current pattern of LDS fires to a pattern of cooler, early dry season (EDS) fires, and that the vast majority of global savanna fire emissions (74%) occur in Africa across 20 least developed countries (LDC). Other studies suggest that similar fire abatement can increase soil and woody carbon sequestration^52,53^.

Here, we investigate the potential for a well-managed EDS fire management program to generate carbon credits for African lion PAs and a new, sustainable financial revenue stream. We demonstrate this financing could indirectly support lion conservation through habitat restoration (by reducing catastrophic fires and associated damage to soils and vegetation) and directly support lion conservation via other PA management actions (e.g., increased anti-poaching efforts and improved human-wildlife conflict resolution). We used greenhouse gas (GHG) emission abatement from methane (CH_4_) and nitrous oxide (N_2_O) to serve as a baseline, a relatively conservative estimate of potential carbon revenue (PCR) that could be generated from an EDS fire program^51^. We then combined emission abatement potential with estimates of carbon sequestration from three other carbon pools (i.e., non-living biomass^54,55^, living biomass^56^, and soil carbon^57^) to produce an upper estimate of carbon credit revenue generating potential. We combined these estimates to produce a range of PCR to determine how much of the lion PA financial shortfall could be met from carbon credits generated by EDS savanna burning program that would support more effective PA management for lions. We discuss limitations of this approach in the context of land degradation trends and the stark realities of economic recovery from COVID-19 in some of the least developed countries on the planet. We conclude by proposing a series of lion-centric pilot projects as symbolic projects to launch the UNs Decade of Ecological Restoration by envisioning savanna fire management programs as natural climate solutions and conservation-development investments that could reap multiple environmental and societal benefits in Africa.

## RESULTS

### Potential carbon revenue from savannah fire management for lion PAs in Africa

PCR from an EDS savanna fire management program could partially or entirely close the estimated US$1B funding shortfall for managing lion PAs in Africa. We found that for all lion PAs in Africa, an EDS fire management program could generate PCR ranging from USD $59.6 - $655.9 M per year (Table 1) based on the lower voluntary market average price (USD $5/ton). At a higher price of USD $13/ton, such a program could generate USD $155.0 M – $1.7 B per year (Table 2).

**Table 1:**
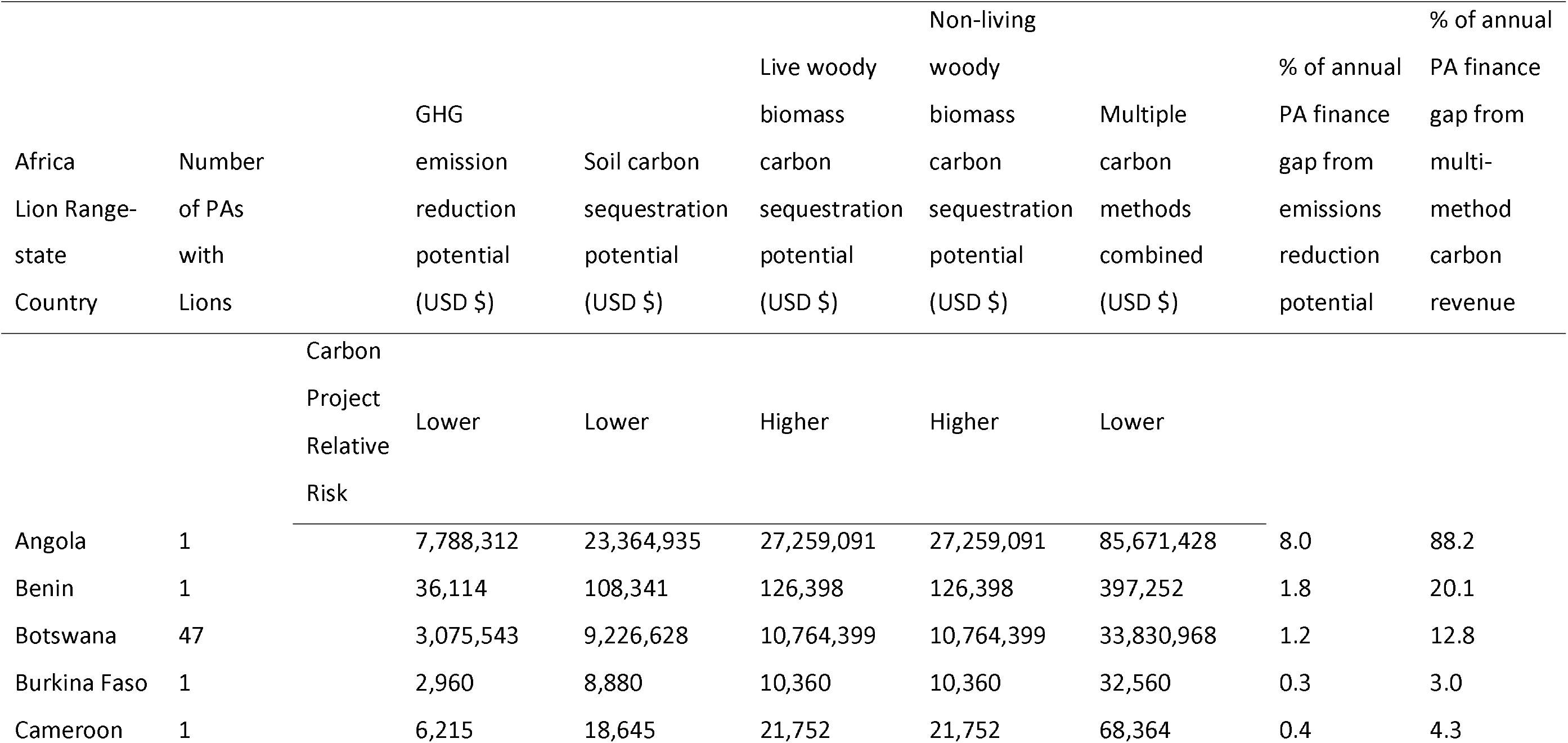
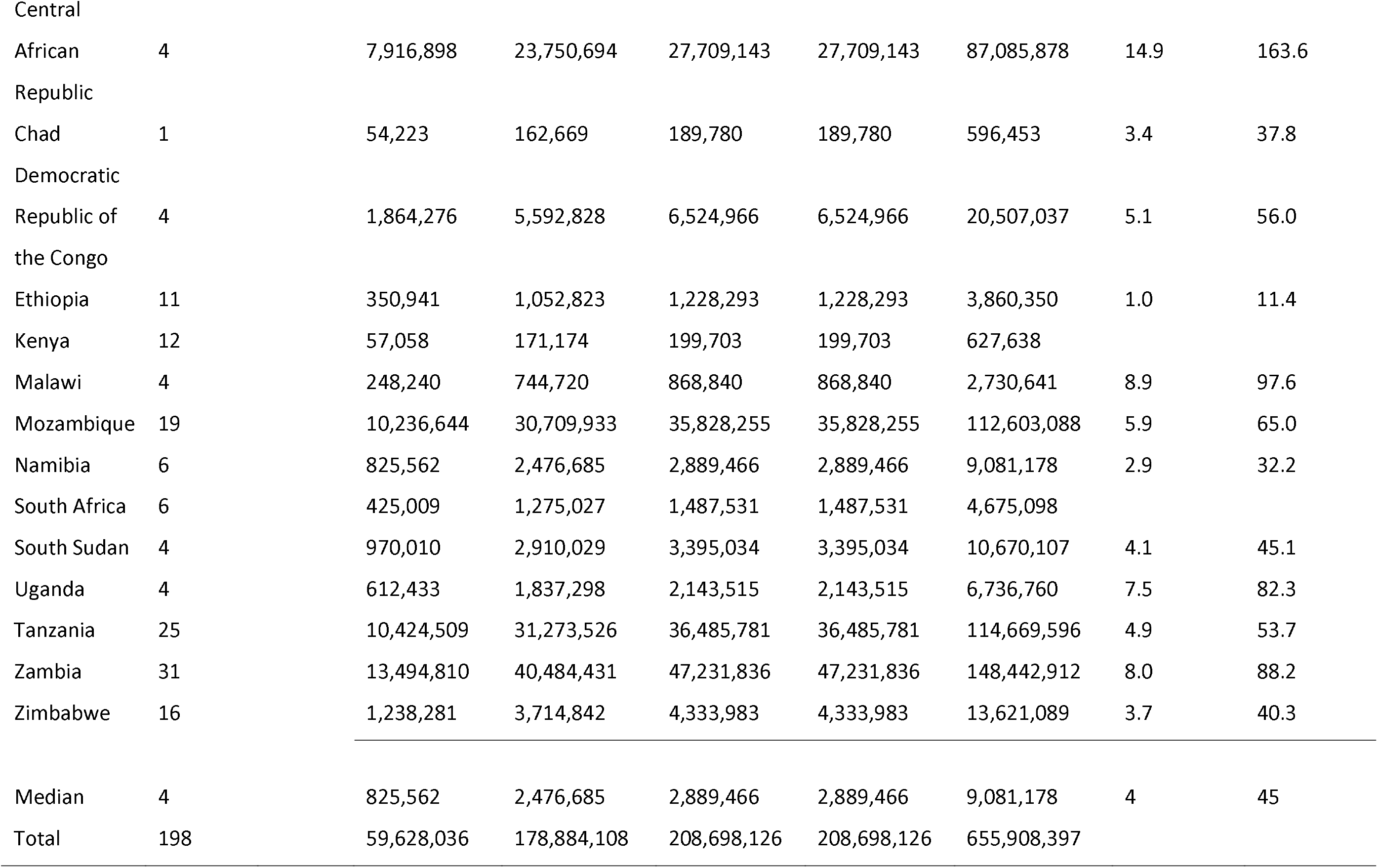
African Lion range states and the potential for EDS fire management programs to generate carbon revenue in relation to protected area (PA) finance gaps for all PAs (with >0 greenhouse gas (GHG) reduction potential). The rate of USD $5/ton was applied as a lower estimate based on the current average of voluntary market values. Multiple methods include four existing carbon methodologies that could generate carbon credits from implementing an EDS fire management program focused on emissions reduction and carbon sequestration. PA finance gaps were based on cost estimates for effective PA management according to Lindsey et al^28^.

**Table 2:**
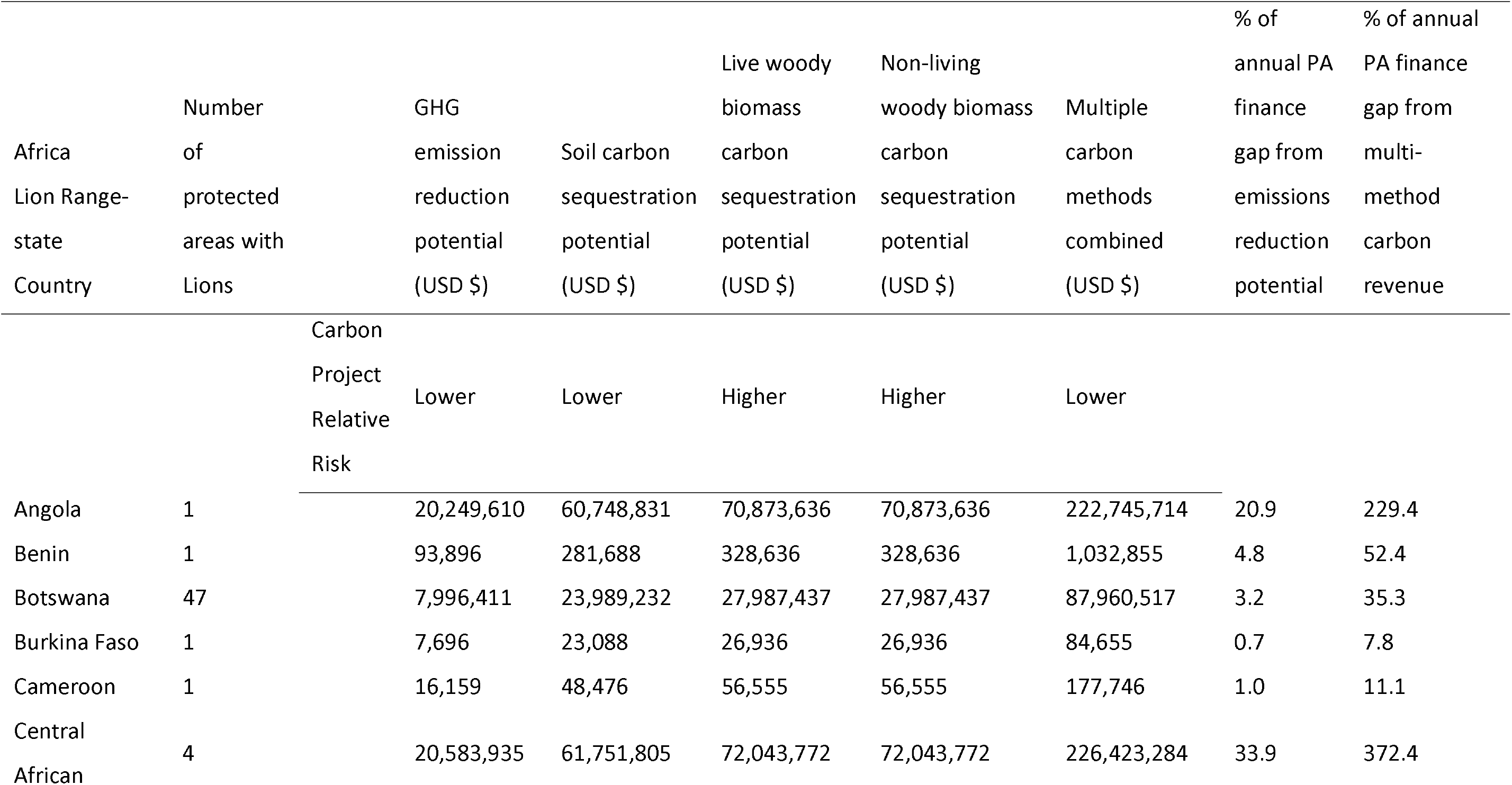
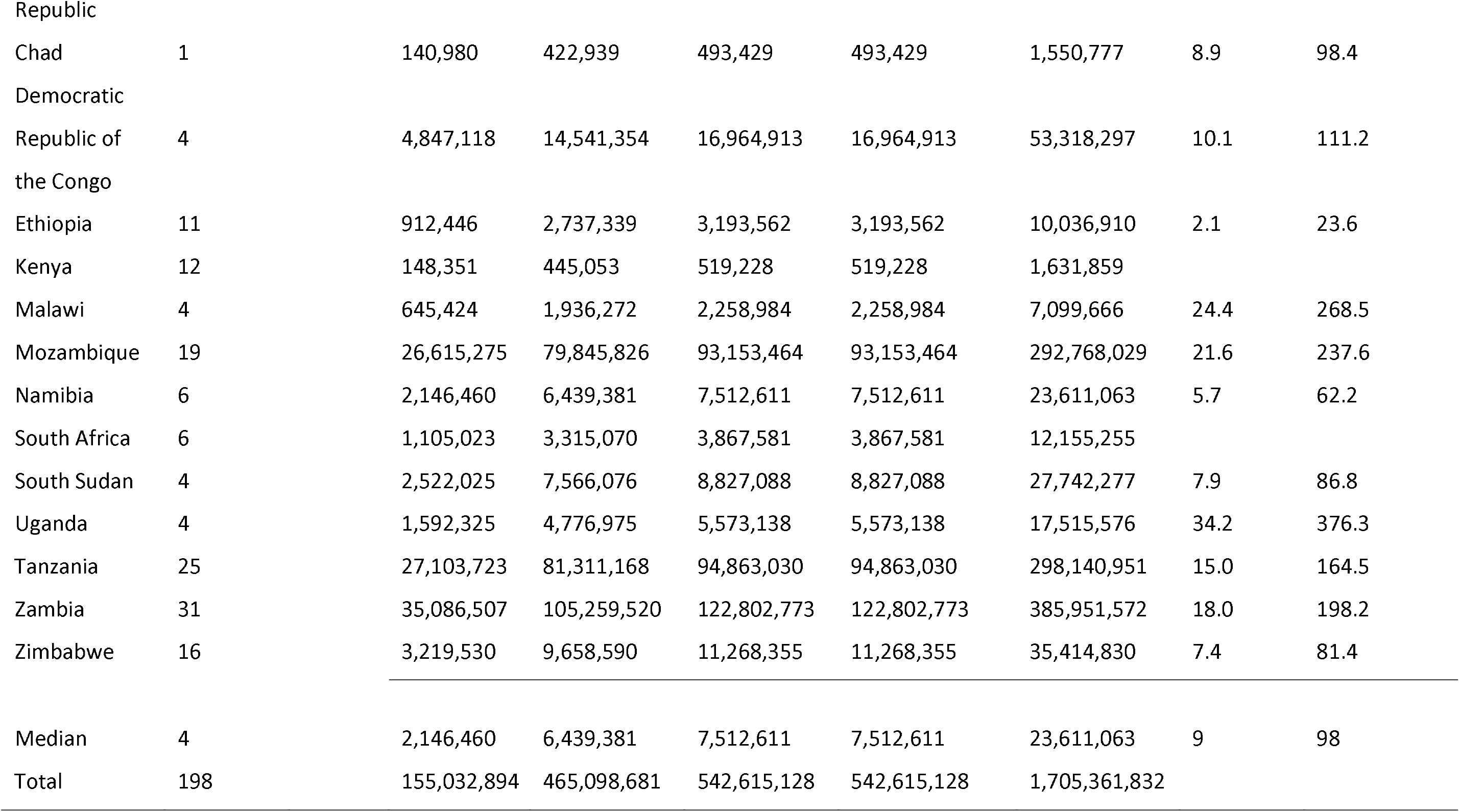
African Lion range states and the potential for EDS fire management programs to generate carbon revenue in relation to protected area (PA) finance gaps for all PAs (with >0 greenhouse gas (GHG) reduction potential). The rate of USD $13/ton was applied as an upper estimate based on an average of Western Climate Initiative Carbon Auction Settlement prices93. Multiple methods include four existing carbon methodologies that could generate carbon credits from implementing an EDS fire management program focused on emissions reduction and carbon sequestration. PA finance gaps were based on cost estimates for effective PA management according to Lindsey et al.^28^.

In countries with the greatest potential for emissions reduction, five of seven (71%) are also countries with the greatest funding needs for lion PA management (Fig. 1). As all seven of the countries with the greatest emissions reduction potential are also characterized as Least Developed Countries (LDCs), there is substantial opportunity for this additional, new funding to provide significant co-benefits to national and local economies.

**Figure 1.**
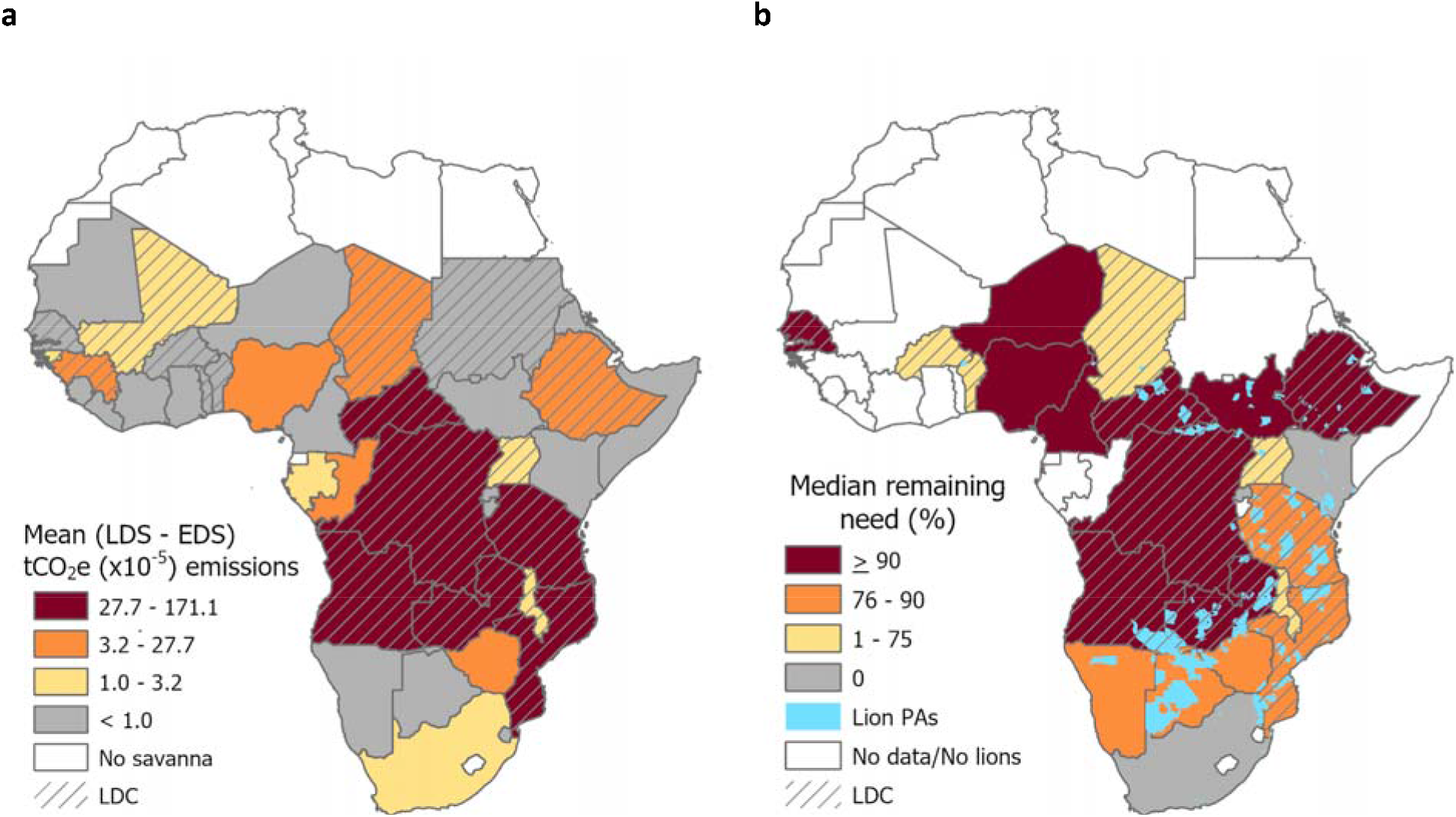
The country-level relationship between a) carbon emission reductions resulting from instituting a fire management plan that shifts fires from the late dry season (LDS) to the early dry season (EDS) compared to b) countries with lions and the financial need estimated for effective protected area (PA) management. The potential benefit of launching EDS savanna burning activities (a) are scaled by mean country level abatement potential (LDS - EDS) emissions (using terciles on GHG values >0), and countries with crosshatching are least developed countries (LDC) (from Lipsett-Moore et al.51).

There is substantial potential for emission reductions benefits for the majority (65%; 198 of 303 in Supplemental Table 1) of all lion PAs (Fig. 2b). There is also substantial variability in the amount of local emissions produced within most landscapes (Fig 2a), and significant trend (R^2^=0.61, p<0.001) that the larger the PA the greater the emissions reduction potential and therefore greater PCR (Fig. 3). The PCR for each lion PA will be a product of the emissions reduction generated and the market price of carbon credits. As would be expected, larger PAs with a history of late dry season (LDS) wildfires and higher productivity generally demonstrate greater revenue generating potential (Supplemental Fig. 1). Emissions reductions alone for all lion PAs across all countries could generate between a median of USD$ 825,562 per country (Table 1 at USD $5/ton) to a median of USD $2.1 M per country (Table 2 at USD $13/ton).

**Figure 2.**
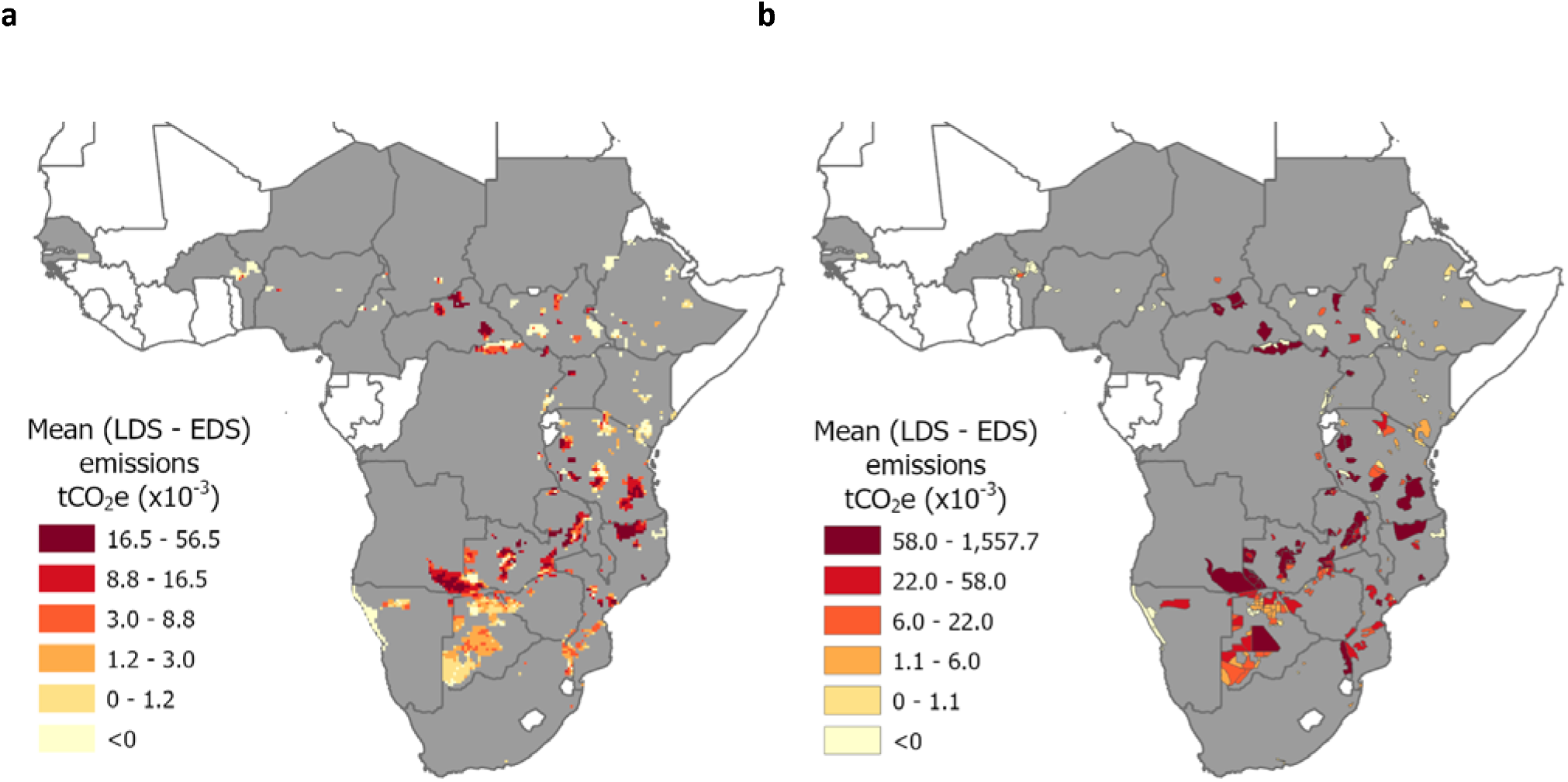
Mean annual emissions abatement potential (LDS-EDS) for 256 protected areas (PA) with lions by a) pixels within lion PAs, and b) summed for the entire lion PA. Classes were based on quintiles (for all values >0). Twenty-three countries (gray) contain at least 1 of the 256 PAs.

**Figure 3.**
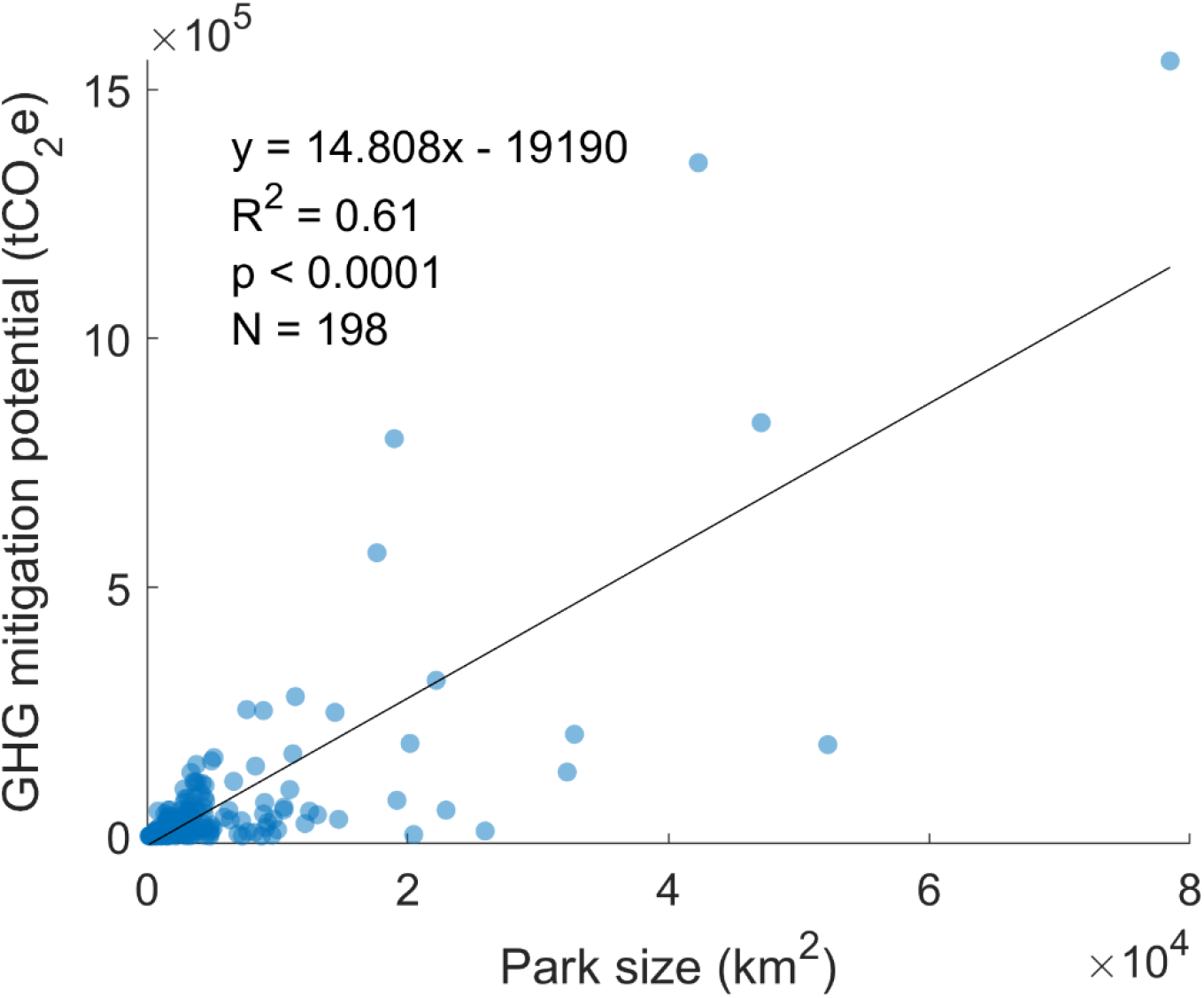
Linear regression showing the significant relationship between protected area size and greenhouse gas emissions reduction potential (>0 tCO_2_-e) from EDS fire management.

However, the same EDS fire management program could also generate additional carbon credits from sequestration across the three remaining carbon pools: (1) non-living biomass, (2) live woody biomass and (3) soil carbon. Our estimates suggest that by shifting to cooler, patchier EDS fires, more PCR would be accrued by combining the carbon sequestration and abatement across all carbon pools (Tables 1 and 2). This would reduce the financial shortfall for each lion range country by 4-9% (median) for emissions abatement alone as compared to 45-98% (median) by combining all carbon pools (at USD $5-13/ton, Tables 1 and 2). While living and non-living biomass PCR are higher than for avoided emissions or soil carbon, they also introduce higher relative risk (Tables 1 and 2). This estimate does not account for other ecosystem services and local employment benefits or costs produced from implementing an EDS fire management program.

### Potential revenue for the top 20 PAs with lions

While there is significant variability in the potential for fire management-generated emissions reductions across all lion PAs in Africa, a fire-based carbon project could have a significant and positive impact on some of the largest and most important lion populations. When we ranked the top 20 PAs prioritized by the greatest PCR, such carbon financing could generate between USD $2.0 M/year on average for each lion PA (at USD $5/ton via emissions abatement only) and USD $57.5 M/year (at USD $13/ton combining all carbon pools) (Table 3). Such funding could potentially restore lion populations and collectively conserve up to 23,191 lions with an average of 1160 lions per PA (Table 3; SD 1196, n=20). Similarly, when we ranked the top twenty lion PAs prioritized by lion population size, their PCRs could potentially support up to 30,563 lions with an average of 1,528 lions per PA (Table 4; SD 1028, n=20), generating on average USD $1.5 M/year per lion PA (at USD $5/ton via emissions reduction) and USD $44.3 M/year (at USD $13/ton via multiple methods) (Table 4).

**Table 3:** Top twenty lion protected areas in Africa estimated to gain the most carbon revenue generated from a fire management program. The low estimate of potential carbon revenue (PCR) was based on emissions reduction (Lipsett-Moore et al. 201851), and valued at an average voluntary carbon market rate of US $5/ton. The higher end of the range added in three carbon sequestration estimates, and was valued at the higher voluntary market average rate of US $13/ton. The middle PCR estimate used multiple methods (emission reduction and sequestration) at the lower market rate. Predicted lion population estimates (i.e., carrying capacity) were created from Loveridge and Canney^67^.

| Rank | Africa<br>Lion Range-state<br>Country | Africa Lion<br>Protected<br>Area | Low PCR/yr<br>\$USD | Middle PCR/yr<br>\$USD | High PCR/yr<br>\$USD | Predicted Lion<br>Population<br>Estimate |
| --- | --- | --- | --- | --- | --- | --- |
| 1 | Angola | Luengue-Luiana | 7,788,312 | 85,671,428 | 222,745,714 | 5,633 |
| 2 | Mozambique | Niassa | 6,765,565 | 74,421,213 | 193,495,153 | 1,881 |
| 3 | Tanzania | Selous | 4,153,721 | 45,690,931 | 118,796,419 | 2,157 |
| 4 | Central African<br>Republic | Manovo-Gounda-Saint Floris | 3,993,089 | 43,923,982 | 114,202,353 | 1,457 |
| 5 | Central African<br>Republic | Chinko | 2,846,294 | 31,309,238 | 81,404,020 | 359 |
| 6 | Zambia | Kafue | 1,564,726 | 17,211,987 | 44,751,166 | 1,714 |
| 7 | Tanzania | Moyowosi | 1,401,352 | 15,414,870 | 40,078,663 | 762 |
| 8 | Zambia | Lunga-Luswishi | 1,271,273 | 13,984,003 | 36,358,407 | 416 |
| 9 | Zambia | South<br>Luangwa | 1,260,617 | 13,866,785 | 36,053,640 | 1,104 |
| 10 | Zambia | Musalangu | 1,242,875 | 13,671,622 | 35,546,217 | 1,594 |
|  | Democratic |  |  |  |  |  |
| 11 | Republic of the | Bili-Uere | 1,022,510 | 11,247,608 | 29,243,781 | 638 |
|  | Congo |  |  |  |  |  |
| 12 | Tanzania | Ruaha | 930,825 | 10,239,071 | 26,621,585 | 540 |
| 13 | Botswana | Central<br>Kalahari | 918,503 | 10,103,529 | 26,269,176 | 1,616 |
| 14 | Central African<br>Republic | Bamingui-Bangoran | 826,465 | 9,091,120 | 23,636,912 | 413 |
| 15 | Tanzania | Ugalla | 788,381 | 8,672,195 | 22,547,707 | 269 |
|  | Democratic |  |  |  |  |  |
| 16 | Republic of the | Garamba | 754,720 | 8,301,918 | 21,584,987 | 95 |
|  | Congo |  |  |  |  |  |
| 17 | Zambia | Kafinda | 719,328 | 7,912,603 | 20,572,767 | 257 |
| 18 | Zambia | Luano | 702,844 | 7,731,285 | 20,101,341 | 800 |
| 19 | Zambia | West<br>Zambezi | 642,913 | 7,072,043 | 18,387,312 | 1,254 |
| 20 | Tanzania | Lwafi | 640,308 | 7,043,384 | 18,312,799 | 231 |
| Mean |  |  | 2,011,731 | 22,129,041 | 57,535,506 | 1,160 |
| SD |  |  | 2,029,972 | 22,329,691 | 58,057,198 | 1,196 |
| Total |  |  | 40,234,620 | 442,580,815 | 1,150,710,118 | 23,191 |

**Table 4:** Top twenty protected areas in Africa with the largest estimated lion populations, and the amount of potential carbon revenue generated from a fire management program. The low estimate of potential carbon revenue (PCR) was based on emissions reduction (Lipsett-Moore et al. 201851), and valued at an average voluntary carbon market rate of US $5/ton. The higher end of the range added in three carbon sequestration estimates, and was valued at the higher voluntary market average rate of US $13/ton. The middle PCR estimate used multiple methods (emission reduction and sequestration) at the lower market rate. Predicted lion population estimates (i.e., carrying capacity) were created from Loveridge and Canney^67^.

| Rank | Africa Lion<br>Range-state Country | Africa Lion<br>Protected Area | Low PCR/yr<br>\$USD | Middle PCR/yr<br>\$USD | High PCR/yr<br>\$USD | Predicted Lion<br>Population Estimate |
| --- | --- | --- | --- | --- | --- | --- |
| 1 | Angola | Luengue-Luiana | 7,788,310 | 85,671,410 | 222,745,666 | 5,633 |
| 2 | Tanzania | Selous | 4,153,720 | 45,690,920 | 118,796,392 | 2,157 |
| 3 | Mozambique | Niassa | 6,765,565 | 74,421,215 | 193,495,159 | 1,881 |
| 4 | South Sudan | Zeraf | 467,260 | 5,139,860 | 13,363,636 | 1,779 |
| 5 | Tanzania | Serengeti | 214,030 | 2,354,330 | 6,121,258 | 1,776 |
| 6 | Zambia | Kafue | 1,564,725 | 17,211,975 | 44,751,135 | 1,715 |
| 7 | Botswana | Central Kalahari | 918,505 | 10,103,555 | 26,269,243 | 1,616 |
| 8 | Zambia | Musalangu | 1,242,875 | 13,671,625 | 35,546,225 | 1,594 |
| 9 | Central African Republic | Manovo-Gounda-Saint Floris | 3,993,090 | 43,923,990 | 114,202,374 | 1,457 |
| 10 | South Sudan | Badingilo | 221,070 | 2,431,770 | 6,322,602 | 1,455 |
| 11 | Zambia | West Zambezi | 642,915 | 7,072,065 | 18,387,369 | 1,254 |
| 12 | Kenya | Tsavo East & West | 14,525 | 159,775 | 415,415 | 1,169 |
| 13 | Zambia | South Luangwa | 1,260,615 | 13,866,765 | 36,053,589 | 1,104 |
| 14 | Ethiopia | Gambella | 226,705 | 2,493,755 | 6,483,763 | 958 |
| 15 | South Africa | Kruger | 359,400 | 3,953,400 | 10,278,840 | 908 |
| 16 | Burkina Faso | Arli | 2,960 | 32,560 | 84,656 | 890 |
| 17 | South Sudan | Meshra | 79,130 | 870,430 | 2,263,118 | 869 |
| 18 | Zambia | Luano | 702,845 | 7,731,295 | 20,101,367 | 800 |
| 19 | Zimbabwe | Hwange | 168,295 | 1,851,245 | 4,813,237 | 779 |
| 20 | Namibia | Etosha | 260,675 | 2,867,425 | 7,455,305 | 769 |
| Mean |  |  | 1,552,361 | 17,075,968 | 44,397,517 | 1,528 |
| SD |  |  | 2,228,922 | 24,518,145 | 63,747,177 | 1,028 |
| Total |  |  | 31,047,215 | 341,519,365 | 887,950,349 | 30,563 |

Given that total lion recovery potential (i.e., carrying capacity) for the 198 lion PAs that could generate fire-carbon revenue is estimated at approximately 60,000 lions (see Supplemental Information – Table 2), prioritizing investments on the 20 lion PAs with the greatest PCR potential would capture one third of the lion recovery potential versus half of recovery potential for the 20 lion PAs with the greatest lion carrying capacities. Furthermore, the top 20 PAs when ranked by PCR (Table 3) occurred in half as many countries (7) as when PAs were ranked by potential lion numbers (14 countries, Table 4). Two countries (Zambia and Tanzania) emerged as having the greatest PCR potential for range-wide lion conservation, as they contained more than half of the lion PAs prioritized by PCR. When the lion PA rankings from both lists were combined (Supplemental Information Table S3), only ten (10) lion PAs occurred on both prioritized lists, of which the top five combined-rank sites each occurred in different countries (illustrating that no country was prioritized). However, Zambia emerged as a clear country-level priority from the complete combined-rank prioritization list, as it occurred multiple times (5), capturing half of all the top 10 priority sites.

## DISCUSSION

Savannas are the world’s most fire prone landscapes, contributing 30% of terrestrial net primary production^58^ and covering 20% of the Earth’s terrestrial surface^59^. Given the global focus on decreasing land degradation to address climate change^40^ paired with greater attention on finding natural climate solutions^48^, our results suggest that fire management could play a critical role in financing conservation of and directly improving savanna habitats. As 20 least developed countries in Africa account for nearly three quarters (74%) of the mitigation potential from an EDS savanna fire management program^31^, there is clearly potential for launching local fire management programs as a bona fide natural climate solution that would directly contribute to national and local economies. This climate-smart strategy is starkly different from many other better-known climate mitigation strategies such as emissions reductions and afforestation in the developed word. Most notably, this brand of natural climate solution intends to generate multiple additional benefits including biodiversity conservation (i.e., lion conservation), ecosystem and economic resilience, and enhanced ecosystem services that will improve human well-being of climate-vulnerable local economies.

Recent analysis of World Bank and Global Environment Facility funding – the largest international donor for biodiversity funding, revealed that integrated conservation and development projects in the tropics did not favor high biodiversity areas, nor countries with the greatest development needs^60^. Furthermore, they found that while Sub Saharan Africa is stated as a top priority for World Bank funding, that African countries have been largely underserved in comparison to other continents in receiving integrated conservation and development project funding. There is great potential for fire management programs to improve local livelihoods by not only hiring local community members, but also acting as a venue to incorporate local ecological knowledge into habitat management programs. Well protected and managed natural habitats help protect key ecosystem services, such as clean water provision, and PAs also have potential to act as hubs for the tourism industry development and other forms of service provision to communities^35^. As the most significant savanna burning occurs in some of the world’s most vulnerable countries, this is an untapped opportunity to invest in integrated conservation-development projects that also contain the vast majority of the world’s remaining free-ranging, but rapidly declining African lion populations^3^.

We acknowledge our predicted lion numbers and PCR estimates could be affected by biases that exaggerate the stated potential. For example, the lion carrying capacity model was based on biological estimates that do not factor in human habitation within and around many of these lion PAs that will reduce potential lion population size regardless of fire programs and within-PA management. However, to estimate the potential for lion recovery, we believe that having a consistent index of lion potential to apply across all lion PAs was more appropriate for this study than accommodating the high uncertainties associated with current lion population estimates^61^. For PCR estimation, current voluntary carbon market rates (USD $2-3/ton) are lower than the 13-year voluntary market average used in this study (USD $5/ton), and we did not include the cost of setting up carbon projects in each of these lion PAs as they are highly variable and difficult to estimate consistently. The current compulsory carbon market rates in Australia average ~USD $7.68/ton and recently sold for USD $10.27/ton, and the introduction of the Carbon Offsetting Scheme for International Aviation (CORSIA), future carbon prices may far exceed those used in this study^49^. Hence, our PCR estimates may be underestimating the carbon values over 5-15 year horizons.

Finally, there is increasing concern that carbon projects in general, and REDD+ projects in particular, carefully assess the measurement, reporting, and verification (MRV) requirements as they can result in prohibitively high costs that ultimately render the project to be financially unsustainable^46^. Unfortunately, as these rangeland carbon projects are so new (both in Africa and for the different carbon methodologies), there is not enough information to model the optimal cost/benefit ratios as has been done for REDD+ forest carbon projects. It is our assumption that larger projects have better cost/benefit ratios in relation to MRV costs than small projects, that in turn must be balanced with sufficient carbon credits from either reduced emissions and/or carbon sequestration. In order to appropriately assess these tradeoffs, more location-specific information is needed, and as this is a continental assessment, it is outside the scope of this assessment. Assuming our estimates represent a credible and appropriate balance between current and future voluntary and compulsory carbon market prices, our analysis confirms that there is significant potential for an effectively run fire management program to provide substantial sources of revenue that would be offset to greater or lesser amounts by the kind of carbon financing chosen, the productivity of the landscape, and the associated MRV costs.

We further recognize that while savanna burning projects in Australia have made a substantial contribution to Australian conservation – generating multiple benefits for PAs and local communities^62^, it will take significant investment for similar benefits to be realized in Africa. A key constraint to the large-scale derivation of carbon revenues in Africa is for African countries to have appropriate regulatory frameworks that guide the generation and reinvestment of carbon revenues in a manner that results in strengthened management of PAs – rather than the funds being captured by the central government. A savanna burning program clearly presents the potential for “new and sustainable financing”, as little carbon financing is currently being generated for lion PAs in Africa. Such projects have been slow to start due to the fact that they reside in savannas, not forests where most of the carbon project investments have been made (though some projects have been started in Kenya, Tanzania, and Zambia). As each of the three carbon sequestration methods introduced in this study are projected to generate revenues for two to three decades^47,54,55^, this type of carbon financing offers a vastly more sustainable alternative to most short-term development aid programs. However, given the multiple socio-cultural and economic benefits of these fire management-based carbon financing projects, we suggest that funding the start-up costs for establishing such a long-term, sustainable financing program could be viewed as a credible, integrated conservation and development project that further supports the global call for ecological restoration.

There are multiple ways such a project can benefit lions in crisis. The revenue generation provided by these carbon projects that is above the costs of implementing the fire management program (and its associated MRV costs) could be used to support other PA lion conservation activities. We recognize that the provision of more money, alone, will not directly result in more lions unless it is provided in the context of sound governance and management structures. For example, surplus resources could be used to support anti-poaching patrols or projects that improve human-wildlife co-existence, such as those delivered by a growing number of effective public-private partnerships for PA management^34^. These activities would further augment the potential for altered fire management to improve lion habitat by increasing forage quality and quantity for prey species. For example, land managers have used prescribed burning in Africa savanna’s to prevent catastrophic fires that negatively impact vegetation (e.g., riparian forests) and infrastructure, and to create benefits for wildlife and livestock including improving forage quality, controlling bush encroachment, reducing tickborne diseases in livestock, and attracting higher densities of grazers^63^. More productive African savannas (based on rainfall and soil nutrients) support greater herbivore biomass, that in turn supports greater carnivore biomass^64–66^, and lion density in PAs is closely related to prey biomass^67^. Furthermore, EDS fires could improve lion fitness by leaving taller grass in the late dry season, as lions hunt more frequently and with greater success with increased cover^68,69^. While such effects are uncertain and difficult to predict^70^, the presence of fire management teams can also become an additional deterrent to poachers by their presence and their ability to remove snares that are an important source of mortality for lions and their prey. More research and monitoring of lion population’s response to any fire management are needed to ensure that management objectives are being achieved.

Despite all these positives, the potential for carbon credits and lion recovery will not be enough to overcome the current challenges to implementation.

First, there must be lasting capacity to implement fire management programs in the form of expertise, governance structures, and tools for management and assessment to effectively implement EDS burning. In Australia, the savanna fire management program has benefited immensely from indigenous knowledge and implementation capacity^71^. In Africa, while similar knowledge exists there needs to be greater investment to support broader, more cohesive governance structures to administer safe and effective burning programs over large areas^12,13,72^. We urge the bilateral and multilateral development and aid sectors to conceptualize this type of project like the UNs Land Degradation Neutrality program describes it – as a way to leverage progress on meeting the Sustainable Development Goals (SDGs) and reducing the biodiversity crisis^73^.

Second, a major challenge is overcoming an under-appreciation of the importance of soil carbon in the fundamental health of ecosystems, adaptation to climate change and as a sink for GHGs^47^. EDS fire management aligns with broader objectives to build healthier soils and restore productive terrestrial ecosystems. Healthier soils capture more carbon, and in turn improve nutrient cycling and water storage capacity^47^. These outcomes have direct benefits for local communities. For example, miombo woodlands in Africa – like those in Niassa - are of global importance for carbon storage and sequestration^74^ and support the daily needs of more than 100 million people^75–77^, yet are likely to experience increased pressure^78–80^ and need more sustainable land management practices^52,76^. Already available carbon methodologies^81^ provide a way to finance and functionally achieve such practices, and much more awareness is needed in the bilateral and multilateral aid sectors to spur greater investment in this kind of nature-based solution.

Third, our results demonstrate a fundamental need for improved technology transfer to develop the full suite of carbon methodologies for EDS burning for Africa. A simplified platform that maps management actions onto complex formal methodologies and that convert actions into ecosystem services such as carbon credits is needed to enable potential carbon project developers to overcome multiple technical tasks in delivering PCRs. These tasks include estimating baselines, potential emissions reduction and sequestration rates, and demonstrating compliance with applicability conditions and additionality for fire management-based carbon projects. Concurrently, there is also the need for technical capacity to track changes in additional socio-cultural co-benefits. In Australia, such tools have greatly enhanced the uptake and development of projects across northern Australia’s savannas resulting in contracts to secure 13.6 million tons CO_2_-e over the next decade^82^. The availability of such tools (e.g., NAFI and SavBAT) for savanna burning carbon projects greatly enhanced savanna burning uptake and adoption^83^.

Despite the significant potential, there are currently no fire management-based carbon projects in Africa. Identifying priority pilot projects will be a key part of moving forward^51^. Our analysis suggests that initiating projects in Zambia could result in a significant return on this investment, as there are substantially more high priority lion carbon projects in Zambia than any other country. Furthermore, only seven countries capture the top 20 potential lion carbon projects, which provides an important subset of socio-political contexts to prioritize initial investments. As there are many socio-political, economic, and bureaucratic issues that would need to be resolved to ensure that in each country, carbon revenues were re-invested into PA management, prioritizing initial investments to demonstrate a proof-of-concept would be essential.

Finding new sources of funding for conservation is often overtaken by other competing societal needs. Lions already have been identified as not only Africa’s most iconic apex predator, but also a link to developing more robust economies in some of the least developed countries on the planet^84^. If current trends are allowed to continue, many African countries will lose their most iconic wildlife species before they have the chance to benefit significantly from them^28^. Our results suggest that investments in natural climate solutions that improve ecosystem health are complementary rather than competitive with other societal priorities. For example, carbon projects may significantly diversify the conservation revenue portfolio. Such diversification might mitigate extensive COVID-19 related economic impacts, including the catastrophic collapse of tourism revenue essential for most of these lion Pas while also building economic resilience to future, unforeseen crises. Carbon projects offer potential “win-win-win” solutions that address the existential threat of climate change, improve the resilience of local communities, and reduce the loss of biodiversity and degradation of land at the same time.

### Concluding Comments

We have shown how a more strategic approach to fire management in Africa has considerable potential to produce a cascade of positive, self-reinforcing impacts for lion PAs and the local communities that rely on them. Exploiting the natural climate solution potential of savannas can unlock sufficient long-term revenue to close budget gaps for enduring and effective management of PAs with lions. That revenue would allow curtailing of the primary short-term drivers of lion and lion-prey declines, particularly poaching and encroachment by people and livestock on PAs. Concomitantly, increased primary productivity arising from changed fire management practices can reverse habitat degradation, build soil carbon and greater resilience to climate change induced drought, and augment the recovery of lion prey species and lions as well. That in turn unlocks the potential for local communities and national governments to benefit from healthy functioning landscapes with thriving wildlife populations.

The mechanism underlying all these impacts also has substantial potential to reduce carbon emissions at a large scale with obvious benefits for the warming climate. Revenues generated from PAs via carbon financing are likely to help offset some of the opportunity costs associated with PAs and increase the political will for the retention of such lands. Africa is facing explosive human population growth, and countries, such as Zambia and Tanzania with above-average proportions of their land area devoted to PAs and high priority lion populations, will experience greater pressure to reallocate land for agriculture and settlement. In this Decade of Ecological Restoration, we suggest restoring EDS fire regimes in African savannas by launching pilot projects in areas with the greatest potential for restoring lion populations and for reducing greenhouse gases. The latter could also satisfy global conservation and development priorities, as these areas also occur in Least Developed Countries with high biodiversity that have until now not received nearly enough support. And without urgent action, this opportunity may soon be lost.

## EXPERIMENTAL PROCEDURES

### Resource Availability

#### Lead Contact

Further information and requests for resources and data should be directed to and will be fulfilled by the lead contact, Timothy H. Tear.

#### Materials availability

This study did not generate new unique materials.

#### Data and code availability

The full dataset for this study is openly available at https://www.dropbox.com/sh/xsq37wmp8o2aieu/AAALH_dqHoXtiYk7a5_5pj_Ja?dl=0

### LDS-EDS GHG estimates

We followed the methods detailed by Lipsett-Moore et al.^51^ that applied a savanna burning approach adopted by the Australian government^81^ and applied them to all savanna habitat globally. To test the applicability of this Australian EDS method in Africa, we used emissions data and its relationship to woody biomass and fire frequency from Niassa National Reserve (Niassa) in Northern Mozambique. At 42,000 km^2^, Niassa is one of Africa’s largest PAs, contains one of the most intact and least disturbed areas of Africa’s deciduous Miombo woodlands in Africa^85^. It occurs within the Eastern Miombo Woodlands terrestrial ecoregion, which is rated as globally outstanding for both its large, intact ecosystems and high biological distinctiveness index, with a conservation status of “relatively stable”^86^ as well as receiving the highest conservation priority category for terrestrial ecoregions across Africa^87^. As Niassa also has some of most extensive information on fire ecology^88^, and contains one of Africa’s largest unfenced lion populations (which until recently was only one of eight populations with more than 1,000 lions^3^), it represents a critical case study for other potential carbon projects. Fifty sampling plots were established and data were collected on vegetation and fire (see methods in Ribeiro et al.^89^). In particular, emissions data were collected by using the IPCC protocol^90^ which estimates emissions as a function of biomass. Woody biomass was estimated by calibrating the ALOS-PALSAR (L band SAR data, 30m spatial resolution), Sentinel 2 (20m spatial resolution) and Landsat 8 (30m spatial resolution) images. Fire frequency data were collected by using a combination of burned area (<u>MCD64A1</u>, 500m spatial resolution) and active fire (<u>MCD14DL</u>, 1km spatial resolution) products from the Moderate Resolution Image Spectoradiometer (MODIS). Linear regression was performed to assess the strength of the relationships between emissions and the presence of fire (fire frequency) and its potential impact on above ground carbon sequestration (live woody biomass) for both the early and late dry seasons (defined by Lipsett-Moore et al.^51^).

### Testing cross-continental model assumptions

To test if the Australian-derived EDS model assumptions are applicable to lion PAs in Africa, we analyzed fire frequency, average emissions, and woody biomass data from Niassa National Reserve in Northern Mozambique, as this PA was one of the highest priority lion PAs in this study (Supplemental Information Table S3), and one of the few with emissions data. In Niassa (Supplemental Information Fig. S1), LDS fires produced significantly higher average emissions and were associated with greater mean number of fires (p<0.001) and less woody biomass (mg/ha) (p<0.02), suggesting LDS fires burned hotter and covered more area and produced greater emissions. In contrast, there were no significant relationships (p>0.05) among EDS emissions or their variability for either fire frequency or woody biomass, suggesting that EDS fires were “cooler” and/or patchy enough that they had weaker, if any effects on woody carbon and emissions. This supports the prediction that LDS fires in Niassa would create more emissions and be more damaging than EDS fires. A fire management program designed to reduce this risk are therefore predicted to generate similar PCR benefits predicted by the Australian EDS approach, however this assumption would need further testing on site following implementation of the desired fire management prescription.

### Carbon credit estimates for multiple methodologies

Significant potential has already been identified to greatly expand the geographic scope of EDS fire management opportunities^91^. The Australian government continues to develop methods that account for the sequestration of the non-living woody carbon pool^92^, and recently approved a new methodology^55^. To estimate the potential for combining these mitigation methodologies from the same EDS fire management program, we relied on estimates from the only existing protocol that has been applied globally – the emissions reduction potential from an EDS fire management program^51^. In Africa, additional carbon credit generating methods have been for avoided emissions in miombo woodlands^81^ and soil carbon^57^.

To estimate the amount of carbon sequestration potential from EDS fire management from the non-living woody carbon pool, we adopted the 3.5x conversion rate from the emission abatement potential recently adopted by the Australian government for high rainfall areas with poor fire histories^55^. We adopted the lower end (3x) of the living woody biomass range (3-4x the emissions abatement potential)^56^. Our soil carbon estimate of 3x abatement potential is derived from data collected in miombo woodlands in Zambia^53^ that suggests significantly higher estimates from altered fire management than global soil carbon estimates for grasslands^47,48^. All of these are broad estimates for the explicit purpose of identifying PCR in this study, and would need to be refined at regional and local scales for use in a carbon project.

### Carbon credit values and relative risk

The range of carbon credit values were based on two general benchmarks. As carbon values vary widely, our lower estimate (USD $5/ton) was based on the average value of carbon credits on the voluntary market over the past 14 years (USD $4.9/ton; 2006-2018, n=13, sd = 1.47)^43^. The higher value (USD $13/ton) was based on Western Climate Initiative Carbon Auction Settlement prices over a similar time frame (USD $13.5/ton; 2012-2018, n=17, sd = 1.12)^93^. Recently, soil carbon credits using a VCS methodology from the Northern Rangelands Trust in Kenya sold for $8/ton (M. Brown pers. comm, Jan 2021), suggesting that our range ($5-$13/ton) reasonably estimates the existing carbon markets. With the rise in interest of nature-based solutions over the past two years, and the increased volume of VCS credits, both the lower and higher estimates are considered conservative estimates as prices are expected to increase in the near future for multiple reasons^43^.

All carbon projects must evaluate risk. While the vast majority of carbon projects focus on forested habitat, grassland carbon projects may produce more reliable carbon sinks than forest carbon projects primarily because catastrophic fires can do more damage to forest carbon stores than to grasslands^94^. Any sequestration project in Australia (including savanna burning projects) have a 25-year or 100-year permanence obligation^55^. In this study, we generally categorized the relative risk of different savanna carbon projects based on the chance that the carbon credits produced by each different methodology could be lost once they were accrued. There is a relatively low risk for activities that produce emissions reduction and soil carbon because they are less likely to be lost than living and non-living woody biomass carbon pools that could be destroyed by catastrophic fires. Combining multiple methods and diversifying strategies was considered a lower relative risk because it contains two sources of lower risk methods.

### Lion PA financial assessments and carrying capacity estimates

The median funding shortfall for lion PA management was assessed for 23 of 27 lion-range countries and for each lion PA based on methods detailed by Lindsey et al.^28^. For this study, we used the middle of three PA area cost estimates (i.e., Lindsey et. al.^28^ method $1,271/km^2^) that was based on modeled costs of managing lion PAs at > 50% of carrying capacity. Given the challenges of consistently and accurately assessing lion population sizes in Africa^61^ we chose to rely on a rangewide model of carrying capacity estimates presented in Lindsey et al.^2^ developed by Loveridge and Canney^67^. Loveridge and Canney’s^67^ biologically-based predictive model was based on published relationships among rainfall, soil nutrient and herbivore biomass in African savannah ecosystems that were combined to predict prey biomass. Then, using prey biomass data from relatively undisturbed and protected sites only, they developed a model to predict lion biomass and density within lion PAs.

## Author Contributions

T.H.T. and G.J.L-M. designed the study. T.H.T., N.H.W., L.S.P., N.S.R., and A.J.L. provided the data analysis. T.H.T., N.H.W., G.J.L-M., M.E.R., N.S.R., L.S.P., P.A.L., L.H., A.J.L., and F.S. interpreted the data and wrote the paper.

## Competing interests

The authors declare no competing interests.

## Additional Information

Supplementary information is available for this paper (three tables and two figures submitted separately).

## Notes

https://www.dropbox.com/sh/xsq37wmp8o2aieu/AAALH_dqHoXtiYk7a5_5pj_Ja?dl=0

### Competing Interest Statement

The authors have declared no competing interest.

