## Supplemental Tables and Figures for "A burning question: Can savannah fire management generate enough carbon revenue to help save the lion from extinction?"

**Supplemental Information**

**Table S1**. Protected areas in Africa with lions by country and identification number (PA ID), the amount of emissions produced from late dry season (LDS) fires (in metric tons (MT) per year and standard deviation (SD)), the mean amount of greenhouse gasses (GHG) avoided in the protected area if fires were shifted to early dry season (EDS) fires, and the associated carbon market values (in USD$ per MT) needed to eliminate the protected area funding deficit from the estimated GHG emissions abatement and combining multiple carbon methodologies.

| Country | PA ID # | Mean  LDS-EDS  MtCO_2_-e yr−1 | SD  LDS-EDS  MtCO_2_-e yr−1 | Mean  LDS-EDS GHG/km^2^ | Mean  LDS-EDS multi/km^2^ | Median deficit/km^2^ | Median  remaining  need % | PAs  with  deficit | Price per  MT of  GHG  Emissions  needed | Price per  MT of  Multiple  Carbon  Methods  needed |
| --- | --- | --- | --- | --- | --- | --- | --- | --- | --- | --- |
| Angola | 7 | 1557662.34 | 308083.68 | 19.85 | 218.32 | 1237 | 97 | 100 | 62.33 | 0.44 |
| Mozambique | 279 | 1353112.96 | 201729.31 | 31.99 | 351.89 | 1136 | 89 | 90 | 35.51 | 0.25 |
| Tanzania | 194 | 830744.19 | 318547.65 | 17.63 | 193.96 | 1095 | 86 | 95 | 62.10 | 0.44 |
| Central African Republic | 57 | 798617.86 | 125870.94 | 42.10 | 463.08 | 1250 | 98 | 75 | 29.69 | 0.21 |
| Central African Republic | 67 | 569258.88 | 157272.80 | 32.27 | 355.02 | 1250 | 98 | 75 | 38.73 | 0.28 |
| Zambia | 221 | 312945.22 | 266639.30 | 14.11 | 155.17 | 1155 | 91 | 100 | 81.88 | 0.59 |
| Tanzania | 196 | 280270.37 | 108328.99 | 24.62 | 270.87 | 1095 | 86 | 95 | 44.47 | 0.32 |
| Zambia | 252 | 254254.59 | 43565.70 | 33.26 | 365.84 | 1155 | 91 | 100 | 34.73 | 0.25 |
| Zambia | 251 | 252123.36 | 36006.63 | 28.23 | 310.52 | 1155 | 91 | 100 | 40.92 | 0.29 |
| Zambia | 238 | 248574.94 | 64273.30 | 17.23 | 189.56 | 1155 | 91 | 100 | 67.02 | 0.48 |
| Democratic Republic of the Congo | 64 | 204501.97 | 182894.98 | 6.24 | 68.60 | 1155 | 91 | 100 | 185.20 | 1.33 |
| Tanzania | 189 | 186164.93 | 59930.28 | 9.23 | 101.48 | 1095 | 86 | 95 | 118.69 | 0.85 |
| Botswana | 1 | 183700.53 | 260622.02 | 3.52 | 38.69 | 1071 | 84 | 100 | 304.48 | 2.17 |
| Central African Republic | 58 | 165293.09 | 160961.04 | 14.77 | 162.47 | 1250 | 98 | 75 | 84.63 | 0.60 |
| Tanzania | 219 | 157676.27 | 79323.29 | 30.66 | 337.30 | 1095 | 86 | 95 | 35.71 | 0.25 |
| Democratic Republic of the Congo | 63 | 150943.97 | 47469.29 | 30.50 | 335.52 | 1155 | 91 | 100 | 37.87 | 0.27 |
| Zambia | 241 | 143865.50 | 12634.76 | 37.90 | 416.85 | 1155 | 91 | 100 | 30.48 | 0.22 |
| Zambia | 234 | 140568.82 | 38736.12 | 16.89 | 185.82 | 1155 | 91 | 100 | 68.37 | 0.49 |
| Zambia | 255 | 128582.60 | 115400.38 | 3.99 | 43.88 | 1155 | 91 | 100 | 289.52 | 2.07 |
| Tanzania | 199 | 128061.53 | 33029.76 | 38.04 | 418.43 | 1095 | 86 | 95 | 28.79 | 0.21 |
| Zambia | 229 | 109969.93 | 45508.84 | 16.54 | 181.95 | 1155 | 91 | 100 | 69.83 | 0.50 |
| Tanzania | 215 | 109275.78 | 21822.38 | 30.50 | 335.46 | 1095 | 86 | 95 | 35.91 | 0.26 |
| Uganda | 165 | 108195.82 | 54101.43 | 27.90 | 306.92 | 853 | 67 | 89 | 30.57 | 0.22 |
| Mozambique | 153 | 107852.65 | 27951.04 | 29.24 | 321.63 | 1136 | 89 | 90 | 38.85 | 0.28 |
| Tanzania | 212 | 105885.37 | 25431.05 | 25.03 | 275.29 | 1095 | 86 | 95 | 43.75 | 0.31 |
| Zambia | 222 | 101692.88 | 32094.16 | 22.84 | 251.23 | 1155 | 91 | 100 | 50.57 | 0.36 |
| Mozambique | 117 | 94580.06 | 120687.90 | 33.30 | 366.33 | 1136 | 89 | 90 | 34.11 | 0.24 |
| South Sudan | 160 | 93451.56 | 72191.18 | 8.53 | 93.87 | 1226 | 96 | 100 | 143.66 | 1.02 |
| Zambia | 245 | 86045.41 | 20743.96 | 20.83 | 229.18 | 1155 | 91 | 100 | 55.44 | 0.40 |
| Zambia | 243 | 76981.64 | 23399.40 | 17.42 | 191.64 | 1155 | 91 | 100 | 66.30 | 0.47 |
| Zambia | 240 | 76542.80 | 12435.95 | 21.22 | 233.46 | 1155 | 91 | 100 | 54.42 | 0.39 |
| Zambia | 253 | 76227.77 | 17846.88 | 24.02 | 264.26 | 1155 | 91 | 100 | 48.08 | 0.34 |
| Zambia | 233 | 75446.18 | 29584.22 | 25.70 | 282.67 | 1155 | 91 | 100 | 44.95 | 0.32 |
| South Africa | 175 | 71880.28 | 69503.57 | 3.75 | 41.24 | 0 | 0 | 22 | 0.00 | 0.00 |
| Zambia | 223 | 69740.95 | 37196.42 | 15.46 | 170.09 | 1155 | 91 | 100 | 74.69 | 0.54 |
| Tanzania | 208 | 67685.82 | 102621.67 | 7.50 | 82.49 | 1095 | 86 | 95 | 146.02 | 1.04 |
| Zambia | 225 | 67504.78 | 7611.83 | 23.65 | 260.14 | 1155 | 91 | 100 | 48.84 | 0.35 |
| Tanzania | 187 | 59245.07 | 33924.94 | 18.20 | 200.15 | 1095 | 86 | 95 | 60.18 | 0.43 |
| Zambia | 254 | 58383.87 | 31276.52 | 13.44 | 147.88 | 1155 | 91 | 100 | 85.91 | 0.62 |
| Botswana | 55 | 58019.83 | 45034.82 | 5.52 | 60.75 | 1071 | 84 | 100 | 193.92 | 1.38 |
| Mozambique | 118 | 52529.34 | 54899.35 | 5.01 | 55.06 | 1136 | 89 | 90 | 226.96 | 1.62 |
| Namibia | 140 | 52246.49 | 24980.18 | 13.60 | 149.60 | 1105 | 87 | 100 | 81.25 | 0.58 |
| Zambia | 248 | 52174.88 | 9878.49 | 32.31 | 355.46 | 1155 | 91 | 100 | 35.74 | 0.26 |
| Namibia | 138 | 52134.84 | 81085.53 | 2.27 | 25.00 | 1105 | 87 | 100 | 486.24 | 3.48 |
| Zambia | 249 | 52079.26 | 19643.21 | 29.55 | 325.00 | 1155 | 91 | 100 | 39.09 | 0.28 |
| Zambia | 231 | 52067.12 | 17705.42 | 16.51 | 181.60 | 1155 | 91 | 100 | 69.96 | 0.50 |
| Namibia | 141 | 51751.44 | 21982.76 | 8.25 | 90.70 | 1105 | 87 | 100 | 134.01 | 0.96 |
| Mozambique | 125 | 51646.23 | 57482.33 | 13.72 | 150.95 | 1136 | 89 | 90 | 82.78 | 0.59 |
| Mozambique | 122 | 51603.11 | 19805.69 | 19.87 | 218.52 | 1136 | 89 | 90 | 57.18 | 0.41 |
| Central African Republic | 56 | 50209.78 | 16504.57 | 59.56 | 655.21 | 1250 | 98 | 75 | 20.99 | 0.15 |
| Botswana | 38 | 50016.38 | 70985.06 | 4.01 | 44.13 | 1071 | 84 | 100 | 266.95 | 1.90 |
| Zambia | 224 | 49928.92 | 19228.77 | 15.75 | 173.26 | 1155 | 91 | 100 | 73.33 | 0.53 |
| Ethiopia | 91 | 45340.84 | 45587.34 | 10.36 | 113.98 | 1208 | 95 | 94 | 116.58 | 0.83 |
| Zambia | 239 | 44246.74 | 7265.99 | 32.44 | 356.80 | 1155 | 91 | 100 | 35.61 | 0.26 |
| South Sudan | 164 | 44213.60 | 49175.11 | 4.95 | 54.41 | 1226 | 96 | 100 | 247.87 | 1.76 |
| Tanzania | 191 | 42805.84 | 91380.41 | 3.28 | 36.03 | 1095 | 86 | 95 | 334.28 | 2.39 |
| Mozambique | 154 | 42791.11 | 30733.24 | 13.47 | 148.19 | 1136 | 89 | 90 | 84.32 | 0.60 |
| Mozambique | 128 | 40796.05 | 47157.74 | 15.03 | 165.30 | 1136 | 89 | 90 | 75.60 | 0.54 |
| South Sudan | 159 | 40510.60 | 16069.90 | 23.17 | 254.85 | 1226 | 96 | 100 | 52.92 | 0.38 |
| Zimbabwe | 266 | 38696.43 | 13151.72 | 11.38 | 125.21 | 1030 | 81 | 100 | 90.49 | 0.65 |
| Mozambique | 149 | 38432.56 | 20553.66 | 10.80 | 118.76 | 1136 | 89 | 90 | 105.22 | 0.75 |
| Zambia | 247 | 38269.01 | 7453.63 | 13.76 | 151.40 | 1155 | 91 | 100 | 83.92 | 0.60 |
| Mozambique | 124 | 37781.71 | 87842.27 | 6.40 | 70.35 | 1136 | 89 | 90 | 177.63 | 1.27 |
| Zambia | 250 | 37367.42 | 12012.90 | 19.50 | 214.48 | 1155 | 91 | 100 | 59.24 | 0.42 |
| Mozambique | 126 | 36305.52 | 15912.62 | 23.33 | 256.59 | 1136 | 89 | 90 | 48.70 | 0.35 |
| Botswana | 284 | 35611.32 | 17337.69 | 12.07 | 132.80 | 1071 | 84 | 100 | 88.71 | 0.63 |
| Botswana | 289 | 34183.60 | 48722.80 | 3.52 | 38.76 | 1071 | 84 | 100 | 303.92 | 2.17 |
| Zimbabwe | 262 | 33658.94 | 32929.91 | 2.29 | 25.19 | 1030 | 81 | 100 | 449.80 | 3.22 |
| Zambia | 246 | 32870.33 | 12250.74 | 21.69 | 238.64 | 1155 | 91 | 100 | 53.24 | 0.38 |
| Mozambique | 120 | 32097.33 | 50692.61 | 5.02 | 55.25 | 1136 | 89 | 90 | 226.16 | 1.61 |
| Mozambique | 150 | 30719.61 | 29819.33 | 4.23 | 46.57 | 1136 | 89 | 90 | 268.32 | 1.91 |
| Malawi | 130 | 30368.97 | 5332.22 | 12.87 | 141.52 | 581 | 46 | 75 | 45.16 | 0.33 |
| Tanzania | 186 | 29811.40 | 13495.83 | 18.80 | 206.84 | 1095 | 86 | 95 | 58.23 | 0.42 |
| Botswana | 39 | 27460.34 | 49103.78 | 2.97 | 32.64 | 1071 | 84 | 100 | 360.92 | 2.57 |
| Botswana | 292 | 25114.62 | 44534.75 | 2.07 | 22.78 | 1071 | 84 | 100 | 517.07 | 3.69 |
| Zambia | 237 | 22971.52 | 18634.55 | 5.26 | 57.82 | 1155 | 91 | 100 | 219.75 | 1.57 |
| Zimbabwe | 270 | 22767.54 | 13861.69 | 7.62 | 83.78 | 1030 | 81 | 100 | 135.24 | 0.97 |
| Zimbabwe | 256 | 22534.22 | 30533.94 | 4.58 | 50.36 | 1030 | 81 | 100 | 224.99 | 1.61 |
| Zimbabwe | 258 | 21958.89 | 7102.78 | 15.44 | 169.81 | 1030 | 81 | 100 | 66.72 | 0.48 |
| Mozambique | 148 | 20919.88 | 10859.71 | 15.16 | 166.76 | 1136 | 89 | 90 | 74.93 | 0.53 |
| Zimbabwe | 257 | 19161.19 | 11164.35 | 9.84 | 108.20 | 1030 | 81 | 100 | 104.71 | 0.75 |
| Zimbabwe | 274 | 18533.25 | 3697.58 | 19.08 | 209.85 | 1030 | 81 | 100 | 53.99 | 0.39 |
| Mozambique | 121 | 18524.75 | 17920.85 | 9.93 | 109.19 | 1136 | 89 | 90 | 114.44 | 0.82 |
| Tanzania | 193 | 17033.97 | 14525.66 | 8.93 | 98.28 | 1095 | 86 | 95 | 122.56 | 0.88 |
| United Republic of Tanzania | 204 | 16969.80 | 99513.97 | 1.86 | 20.50 | 1095 | 86 | 95 | 587.65 | 4.20 |
| Zimbabwe | 265 | 16800.48 | 11369.08 | 9.73 | 106.98 | 1030 | 81 | 100 | 105.91 | 0.76 |
| Ethiopia | 87 | 16322.70 | 7057.49 | 12.78 | 140.54 | 1208 | 95 | 94 | 94.55 | 0.68 |
| Zimbabwe | 272 | 15899.25 | 7908.82 | 7.49 | 82.41 | 1030 | 81 | 100 | 137.48 | 0.98 |
| South Sudan | 157 | 15826.17 | 4400.35 | 3.58 | 39.41 | 1226 | 96 | 100 | 342.19 | 2.44 |
| Mozambique | 151 | 15764.92 | 10596.54 | 3.85 | 42.38 | 1136 | 89 | 90 | 294.84 | 2.10 |
| Botswana | 41 | 15624.59 | 21811.21 | 3.27 | 35.93 | 1071 | 84 | 100 | 327.92 | 2.34 |
| Tanzania | 197 | 15504.64 | 30166.54 | 3.06 | 33.63 | 1095 | 86 | 95 | 358.16 | 2.56 |
| Zimbabwe | 268 | 14741.05 | 7133.03 | 5.10 | 56.15 | 1030 | 81 | 100 | 201.78 | 1.44 |
| Botswana | 8 | 14082.11 | 14610.76 | 6.19 | 68.09 | 1071 | 84 | 100 | 173.03 | 1.23 |
| Botswana | 288 | 13732.60 | 16929.23 | 5.22 | 57.40 | 1071 | 84 | 100 | 205.24 | 1.46 |
| Zambia | 235 | 13607.89 | 5444.55 | 17.81 | 195.96 | 1155 | 91 | 100 | 64.83 | 0.46 |
| Botswana | 17 | 13514.25 | 17079.01 | 3.65 | 40.20 | 1071 | 84 | 100 | 293.04 | 2.09 |
| Mozambique | 119 | 13062.39 | 6084.85 | 4.09 | 44.95 | 1136 | 89 | 90 | 277.98 | 1.98 |
| Botswana | 44 | 12865.66 | 31842.79 | 1.29 | 14.14 | 1071 | 84 | 100 | 833.23 | 5.94 |
| Democratic Republic of the Congo | 283 | 11201.76 | 10409.70 | 7.63 | 83.95 | 1155 | 91 | 100 | 151.35 | 1.08 |
| Chad | 96 | 10844.59 | 13537.23 | 3.56 | 39.21 | 518 | 41 | 100 | 145.34 | 1.05 |
| Tanzania | 210 | 10263.02 | 4532.99 | 16.98 | 186.80 | 1095 | 86 | 95 | 64.48 | 0.46 |
| Botswana | 3 | 10052.58 | 22776.22 | 0.39 | 4.26 | 1071 | 84 | 100 | 2763.53 | 19.70 |
| Malawi | 132 | 9773.63 | 9056.01 | 13.87 | 152.55 | 581 | 46 | 75 | 41.89 | 0.30 |
| Botswana | 6 | 9699.62 | 20541.07 | 1.98 | 21.81 | 1071 | 84 | 100 | 540.16 | 3.85 |
| Botswana | 11 | 9679.62 | 11568.84 | 6.49 | 71.44 | 1071 | 84 | 100 | 164.90 | 1.18 |
| Zambia | 232 | 9643.31 | 35374.34 | 2.55 | 28.03 | 1155 | 91 | 100 | 453.30 | 3.25 |
| Uganda | 168 | 9578.42 | 5818.85 | 14.19 | 156.05 | 853 | 67 | 89 | 60.13 | 0.43 |
| Botswana | 4 | 9534.00 | 17891.73 | 3.76 | 41.33 | 1071 | 84 | 100 | 285.02 | 2.03 |
| Botswana | 48 | 8534.81 | 29100.13 | 1.10 | 12.15 | 1071 | 84 | 100 | 969.44 | 6.91 |
| Zambia | 242 | 8194.89 | 10632.78 | 10.28 | 113.04 | 1155 | 91 | 100 | 112.39 | 0.80 |
| Zimbabwe | 264 | 8181.67 | 6415.79 | 4.79 | 52.65 | 1030 | 81 | 100 | 215.21 | 1.54 |
| Namibia | 142 | 7734.07 | 2602.51 | 10.80 | 118.85 | 1105 | 87 | 100 | 102.27 | 0.73 |
| Tanzania | 183 | 7315.75 | 5132.76 | 0.89 | 9.75 | 1095 | 86 | 95 | 1234.87 | 8.82 |
| Benin | 53 | 7222.77 | 18963.29 | 2.62 | 28.77 | 714 | 56 | 100 | 273.00 | 1.95 |
| Botswana | 294 | 6627.43 | 15733.79 | 2.02 | 22.25 | 1071 | 84 | 100 | 529.42 | 3.77 |
| Zimbabwe | 269 | 6305.64 | 1821.02 | 12.66 | 139.29 | 1030 | 81 | 100 | 81.34 | 0.58 |
| Democratic Republic of the Congo | 65 | 6207.52 | 27109.15 | 2.70 | 29.65 | 1155 | 91 | 100 | 428.51 | 3.07 |
| Botswana | 13 | 5984.10 | 13592.52 | 3.42 | 37.64 | 1071 | 84 | 100 | 312.99 | 2.23 |
| Ethiopia | 89 | 5955.41 | 8733.10 | 2.58 | 28.39 | 1208 | 95 | 94 | 468.10 | 3.35 |
| Botswana | 37 | 5939.05 | 12432.18 | 2.13 | 23.43 | 1071 | 84 | 100 | 502.88 | 3.59 |
| Tanzania | 190 | 5621.95 | 3651.52 | 4.92 | 54.14 | 1095 | 86 | 95 | 222.49 | 1.59 |
| Botswana | 293 | 5519.18 | 11512.35 | 2.19 | 24.11 | 1071 | 84 | 100 | 488.73 | 3.48 |
| South Africa | 173 | 5515.42 | 5581.39 | 9.36 | 102.93 | 0 | 0 | 22 | 0.00 | 0.00 |
| Zimbabwe | 267 | 5485.56 | 3029.58 | 4.60 | 50.55 | 1030 | 81 | 100 | 224.15 | 1.60 |
| South Africa | 220 | 5390.49 | 7105.53 | 5.78 | 63.55 | 0 | 0 | 22 | 0.00 | 0.00 |
| Tanzania | 192 | 5304.00 | 8074.10 | 1.95 | 21.43 | 1095 | 86 | 95 | 561.97 | 4.01 |
| Tanzania | 206 | 5216.56 | 30717.51 | 1.77 | 19.51 | 1095 | 86 | 95 | 617.33 | 4.41 |
| Botswana | 2 | 5179.19 | 8510.43 | 2.00 | 22.00 | 1071 | 84 | 100 | 535.45 | 3.82 |
| Malawi | 131 | 5009.72 | 1100.34 | 9.91 | 109.06 | 581 | 46 | 75 | 58.60 | 0.42 |
| Botswana | 10 | 4915.81 | 7041.70 | 3.66 | 40.24 | 1071 | 84 | 100 | 292.76 | 2.09 |
| Botswana | 16 | 4774.16 | 9283.27 | 1.25 | 13.79 | 1071 | 84 | 100 | 854.51 | 6.09 |
| Uganda | 178 | 4523.56 | 4909.29 | 8.43 | 92.69 | 853 | 67 | 89 | 101.23 | 0.72 |
| Malawi | 133 | 4495.69 | 3515.62 | 4.58 | 50.39 | 581 | 46 | 75 | 126.83 | 0.91 |
| Botswana | 5 | 4478.38 | 6865.00 | 0.93 | 10.20 | 1071 | 84 | 100 | 1154.48 | 8.23 |
| Mozambique | 152 | 4428.00 | 7662.14 | 6.59 | 72.46 | 1136 | 89 | 90 | 172.46 | 1.23 |
| Mozambique | 129 | 4380.69 | 6916.11 | 4.21 | 46.35 | 1136 | 89 | 90 | 269.61 | 1.92 |
| Botswana | 45 | 4367.23 | 13125.14 | 0.63 | 6.89 | 1071 | 84 | 100 | 1710.44 | 12.20 |
| Botswana | 18 | 4353.68 | 9480.28 | 2.00 | 21.95 | 1071 | 84 | 100 | 536.72 | 3.83 |
| Botswana | 40 | 4323.63 | 7150.51 | 3.83 | 42.08 | 1071 | 84 | 100 | 279.96 | 2.00 |
| Zambia | 230 | 4256.04 | 14380.51 | 2.37 | 26.07 | 1155 | 91 | 100 | 487.32 | 3.49 |
| Botswana | 12 | 3558.62 | 6681.53 | 2.15 | 23.69 | 1071 | 84 | 100 | 497.31 | 3.55 |
| Kenya | 99 | 3502.78 | 6054.58 | 1.98 | 21.82 | 0 | 0 | 30 | 0.00 | 0.00 |
| Botswana | 35 | 3287.44 | 4987.18 | 2.92 | 32.14 | 1071 | 84 | 100 | 366.53 | 2.61 |
| Botswana | 19 | 3128.15 | 5184.81 | 1.08 | 11.86 | 1071 | 84 | 100 | 993.10 | 7.08 |
| Kenya | 103 | 2905.13 | 5912.12 | 0.14 | 1.56 | 0 | 0 | 30 | 0.00 | 0.00 |
| Botswana | 26 | 2734.33 | 5235.67 | 0.79 | 8.73 | 1071 | 84 | 100 | 1349.54 | 9.62 |
| Zimbabwe | 271 | 2658.75 | 3203.00 | 5.16 | 56.74 | 1030 | 81 | 100 | 199.69 | 1.43 |
| Kenya | 102 | 2429.78 | 1759.02 | 3.31 | 36.45 | 0 | 0 | 30 | 0.00 | 0.00 |
| Botswana | 20 | 2214.33 | 4787.38 | 1.05 | 11.55 | 1071 | 84 | 100 | 1020.30 | 7.27 |
| Ethiopia | 92 | 2085.42 | 6059.21 | 1.12 | 12.29 | 1208 | 95 | 94 | 1081.54 | 7.73 |
| Botswana | 32 | 2058.18 | 3301.13 | 1.22 | 13.44 | 1071 | 84 | 100 | 876.63 | 6.25 |
| Zambia | 236 | 1833.45 | 12315.79 | 1.56 | 17.14 | 1155 | 91 | 100 | 741.26 | 5.31 |
| Botswana | 25 | 1806.35 | 2792.65 | 2.10 | 23.15 | 1071 | 84 | 100 | 508.83 | 3.63 |
| Tanzania | 198 | 1794.75 | 1008.61 | 2.02 | 22.24 | 1095 | 86 | 95 | 541.58 | 3.87 |
| Botswana | 42 | 1655.68 | 3588.68 | 0.98 | 10.80 | 1071 | 84 | 100 | 1090.68 | 7.78 |
| South Africa | 172 | 1530.85 | 4056.29 | 0.16 | 1.76 | 0 | 0 | 22 | 0.00 | 0.00 |
| Cameroon | 62 | 1242.98 | 2341.18 | 0.88 | 9.72 | 1143 | 90 | 100 | 1293.89 | 9.26 |
| Namibia | 139 | 1087.08 | 972.95 | 3.22 | 35.41 | 1105 | 87 | 100 | 343.28 | 2.46 |
| Kenya | 104 | 1002.34 | 1199.05 | 1.14 | 12.55 | 0 | 0 | 30 | 0.00 | 0.00 |
| Tanzania | 200 | 976.94 | 8106.60 | 0.22 | 2.39 | 1095 | 86 | 95 | 5034.68 | 35.95 |
| Botswana | 43 | 956.74 | 1963.90 | 1.40 | 15.41 | 1071 | 84 | 100 | 764.63 | 5.45 |
| Botswana | 9 | 936.43 | 1368.54 | 0.77 | 8.52 | 1071 | 84 | 100 | 1382.13 | 9.85 |
| Botswana | 291 | 877.17 | 1449.34 | 3.13 | 34.48 | 1071 | 84 | 100 | 341.65 | 2.44 |
| Kenya | 105 | 842.03 | 999.56 | 0.66 | 7.26 | 0 | 0 | 30 | 0.00 | 0.00 |
| Botswana | 290 | 840.51 | 2086.04 | 0.73 | 8.04 | 1071 | 84 | 100 | 1464.94 | 10.45 |
| Botswana | 46 | 678.67 | 1549.72 | 1.36 | 14.91 | 1071 | 84 | 100 | 790.22 | 5.63 |
| Tanzania | 216 | 602.32 | 720.25 | 0.79 | 8.66 | 1095 | 86 | 95 | 1390.70 | 9.93 |
| Burkina Faso | 79 | 591.99 | 7147.29 | 0.49 | 5.41 | 901 | 71 | 100 | 1832.59 | 13.13 |
| Botswana | 285 | 585.05 | 1327.57 | 0.66 | 7.21 | 1071 | 84 | 100 | 1633.44 | 11.65 |
| Botswana | 22 | 574.80 | 1426.04 | 3.37 | 37.10 | 1071 | 84 | 100 | 317.52 | 2.26 |
| Botswana | 286 | 543.63 | 1546.16 | 0.92 | 10.17 | 1071 | 84 | 100 | 1158.15 | 8.26 |
| Kenya | 110 | 490.15 | 4022.13 | 0.56 | 6.16 | 0 | 0 | 30 | 0.00 | 0.00 |
| Botswana | 36 | 417.59 | 1138.46 | 0.14 | 1.55 | 1071 | 84 | 100 | 7623.68 | 54.36 |
| Tanzania | 185 | 410.80 | 1225.33 | 0.25 | 2.73 | 1095 | 86 | 95 | 4404.45 | 31.45 |
| South Africa | 181 | 399.92 | 603.14 | 0.80 | 8.75 | 0 | 0 | 22 | 0.00 | 0.00 |
| South Africa | 176 | 284.84 | 918.32 | 0.26 | 2.83 | 0 | 0 | 22 | 0.00 | 0.00 |
| Botswana | 47 | 249.46 | 2380.52 | 0.67 | 7.42 | 1071 | 84 | 100 | 1587.35 | 11.32 |
| Tanzania | 188 | 236.46 | 2989.11 | 0.07 | 0.78 | 1095 | 86 | 95 | 15358.12 | 109.66 |
| Uganda | 169 | 188.74 | 513.98 | 0.52 | 5.67 | 853 | 67 | 89 | 1655.83 | 11.82 |
| Zimbabwe | 276 | 170.14 | 443.32 | 0.99 | 10.94 | 1030 | 81 | 100 | 1035.56 | 7.40 |
| Namibia | 134 | 158.41 | 528.26 | 0.61 | 6.74 | 1105 | 87 | 100 | 1802.39 | 12.90 |
| Ethiopia | 95 | 126.46 | 449.54 | 0.35 | 3.83 | 1208 | 95 | 94 | 3467.36 | 24.79 |
| Botswana | 21 | 117.10 | 898.81 | 0.51 | 5.61 | 1071 | 84 | 100 | 2101.19 | 14.98 |
| Kenya | 106 | 109.51 | 499.94 | 0.17 | 1.92 | 0 | 0 | 30 | 0.00 | 0.00 |
| Zimbabwe | 261 | 103.15 | 204.23 | 0.10 | 1.05 | 1030 | 81 | 100 | 10828.17 | 77.41 |
| Ethiopia | 94 | 99.80 | 148.81 | 0.01 | 0.12 | 1208 | 95 | 94 | 106332.02 | 760.20 |
| Ethiopia | 85 | 77.97 | 160.42 | 0.23 | 2.51 | 1208 | 95 | 94 | 5304.26 | 37.92 |
| Kenya | 304 | 67.56 | 186.70 | 0.04 | 0.49 | 0 | 0 | 30 | 0.00 | 0.00 |
| Ethiopia | 84 | 66.30 | 92.84 | 0.03 | 0.33 | 1208 | 95 | 94 | 40061.15 | 286.41 |
| Ethiopia | 280 | 50.13 | 200.92 | 0.01 | 0.08 | 1208 | 95 | 94 | 176108.28 | 1259.05 |
| Kenya | 115 | 43.32 | 135.04 | 0.07 | 0.76 | 0 | 0 | 30 | 0.00 | 0.00 |
| Ethiopia | 281 | 38.11 | 164.78 | 0.02 | 0.27 | 1208 | 95 | 94 | 49150.78 | 351.39 |
| Ethiopia | 93 | 25.02 | 135.78 | 0.01 | 0.06 | 1208 | 95 | 94 | 233270.32 | 1667.72 |
| Tanzania | 214 | 24.23 | 7408.24 | 0.03 | 0.36 | 1095 | 86 | 95 | 33041.83 | 235.92 |
| Kenya | 113 | 17.82 | 114.04 | 0.06 | 0.64 | 0 | 0 | 30 | 0.00 | 0.00 |
| Kenya | 101 | 0.84 | 3.29 | 0.00 | 0.02 | 0 | 0 | 30 | 0.00 | 0.00 |
| Kenya | 112 | 0.35 | 1.28 | 0.00 | 0.02 | 0 | 0 | 30 | 0.00 | 0.00 |

**Table S2**. Protected areas (PA) in Africa with lions ranked by potential carbon revenue (PCR) with PA name and type and estimates for low, middle, and high revenue potential (this study), and the predicted number of lions.

| Rank by Revenue | Country | Lion PA Name | PA Type | LDS-EDS emission reduction potential carbon revenue/yr $USD (Low) | Multi-methodology potential carbon revenue/yr $USD (Middle) | Multi-methodology potential carbon revenue/yr $USD (High) | Predicted number of Lions |
| --- | --- | --- | --- | --- | --- | --- | --- |
| 1 | Angola | Luengue-Luiana | National park | 7,788,310 | 85,671,410 | 222,745,666 | 5,633 |
| 2 | Mozambique | Niassa | Game reserve / National reserve / National sanctuary | 6,765,565 | 74,421,215 | 193,495,159 | 1,881 |
| 3 | Tanzania | Selous | Game reserve / National reserve / National sanctuary | 4,153,720 | 45,690,920 | 118,796,392 | 2,157 |
| 4 | Central African Republic | Manovo-Gounda-Saint Floris | National park | 3,993,090 | 43,923,990 | 114,202,374 | 1,457 |
| 5 | Central African Republic | Chinko | Wildlife Sanctuary/Reserve | 2,846,295 | 31,309,245 | 81,404,037 | 359 |
| 6 | Zambia | Kafue | National park | 1,564,725 | 17,211,975 | 44,751,135 | 1,715 |
| 7 | Tanzania | Moyowosi | Game reserve / National reserve / National sanctuary | 1,401,350 | 15,414,850 | 40,078,610 | 762 |
| 8 | Zambia | Lunga-Luswishi | GCA/ WMA / GMA/ CHA | 1,271,275 | 13,984,025 | 36,358,465 | 416 |
| 9 | Zambia | South Luangwa | National park | 1,260,615 | 13,866,765 | 36,053,589 | 1,104 |
| 10 | Zambia | Musalangu | GCA/ WMA / GMA/ CHA | 1,242,875 | 13,671,625 | 35,546,225 | 1,594 |
| 11 | Democratic Republic of the Congo | Bili-Uere | GCA/ WMA / GMA/ CHA | 1,022,510 | 11,247,610 | 29,243,786 | 638 |
| 12 | Tanzania | Ruaha | National park | 930,825 | 10,239,075 | 26,621,595 | 540 |
| 13 | Botswana | CKGR (Central Kalahari) | Game reserve / National reserve / National sanctuary | 918,505 | 10,103,555 | 26,269,243 | 1,616 |
| 14 | Central African Republic | Bamingui-Bangoran | National park | 826,465 | 9,091,115 | 23,636,899 | 413 |
| 15 | Tanzania | Ugalla | Game reserve / National reserve / National sanctuary | 788,380 | 8,672,180 | 22,547,668 | 269 |
| 16 | Democratic Republic of the Congo | Garamba | National park | 754,720 | 8,301,920 | 21,584,992 | 95 |
| 17 | Zambia | Kafinda | GCA/ WMA / GMA/ CHA | 719,330 | 7,912,630 | 20,572,838 | 257 |
| 18 | Zambia | Luano | GCA/ WMA / GMA/ CHA | 702,845 | 7,731,295 | 20,101,367 | 800 |
| 19 | Zambia | West Zambezi | GCA/ WMA / GMA/ CHA | 642,915 | 7,072,065 | 18,387,369 | 1,254 |
| 20 | Tanzania | Lwafi | Game reserve / National reserve / National sanctuary | 640,310 | 7,043,410 | 18,312,866 | 231 |
| 21 | Zambia | Kansonso-Busanga | GCA/ WMA / GMA/ CHA | 549,850 | 6,048,350 | 15,725,710 | 361 |
| 22 | United Republic of Tanzania | Lukwati | Game reserve / National reserve / National sanctuary | 546,380 | 6,010,180 | 15,626,468 | 254 |
| 23 | Uganda | Murchison Falls | National park | 540,980 | 5,950,780 | 15,472,028 | 303 |
| 24 | Mozambique | Gorongosa | National park | 539,265 | 5,931,915 | 15,422,979 | 491 |
| 25 | Tanzania | Rukwa | Game reserve / National reserve / National sanctuary | 529,425 | 5,823,675 | 15,141,555 | 267 |
| 26 | Zambia | Sioma Ngwezi | National park | 508,465 | 5,593,115 | 14,542,099 | 263 |
| 27 | Mozambique | Gil?? | Game reserve / National reserve / National sanctuary | 472,900 | 5,201,900 | 13,524,940 | 74 |
| 28 | South Sudan | Zeraf | Game reserve / National reserve / National sanctuary | 467,260 | 5,139,860 | 13,363,636 | 1,779 |
| 29 | Zambia | Lower Zambezi | National park | 430,225 | 4,732,475 | 12,304,435 | 311 |
| 30 | Zambia | Kaputa | GCA/ WMA / GMA/ CHA | 384,910 | 4,234,010 | 11,008,426 | 250 |
| 31 | Zambia | Munyamadzi | GCA/ WMA / GMA/ CHA | 382,715 | 4,209,865 | 10,945,649 | 445 |
| 32 | Zambia | West Petauke | GCA/ WMA / GMA/ CHA | 381,140 | 4,192,540 | 10,900,604 | 318 |
| 33 | Zambia | Mumbwa | GCA/ WMA / GMA/ CHA | 377,230 | 4,149,530 | 10,788,778 | 185 |
| 34 | South Africa | Kruger National Park | National park | 359,400 | 3,953,400 | 10,278,840 | 908 |
| 35 | Zambia | North Luangwa | National park | 348,705 | 3,835,755 | 9,972,963 | 545 |
| 36 | Tanzania | Kigosi | Game reserve / National reserve / National sanctuary | 338,430 | 3,722,730 | 9,679,098 | 575 |
| 37 | Zambia | Lukusuzi | National park | 337,525 | 3,712,775 | 9,653,215 | 226 |
| 38 | Tanzania | Mikumi | National park | 296,225 | 3,258,475 | 8,472,035 | 501 |
| 39 | Zambia | Lupande | GCA/ WMA / GMA/ CHA | 291,920 | 3,211,120 | 8,348,912 | 503 |
| 40 | Botswana | Chobe | National park | 290,100 | 3,191,100 | 8,296,860 | 705 |
| 41 | Mozambique | Limpopo | National park | 262,645 | 2,889,095 | 7,511,647 | 440 |
| 42 | Namibia | Khaudum | National park | 261,230 | 2,873,530 | 7,471,178 | 198 |
| 43 | Zambia | Lavushi Manda | National park | 260,875 | 2,869,625 | 7,461,025 | 55 |
| 44 | Namibia | Etosha | National park | 260,675 | 2,867,425 | 7,455,305 | 769 |
| 45 | Zambia | West Lunga | National park | 260,395 | 2,864,345 | 7,447,297 | 282 |
| 46 | Zambia | Mulobezi | GCA/ WMA / GMA/ CHA | 260,335 | 2,863,685 | 7,445,581 | 225 |
| 47 | Namibia | Bwabwata | National park | 258,755 | 2,846,305 | 7,400,393 | 346 |
| 48 | Mozambique | Coutada Oficial No. 9 | GCA/ WMA / GMA/ CHA | 258,230 | 2,840,530 | 7,385,378 | 225 |
| 49 | Mozambique | Coutada Oficial No. 10 | GCA/ WMA / GMA/ CHA | 258,015 | 2,838,165 | 7,379,229 | 172 |
| 50 | Central African Republic | Vassako-Bolo | Wildlife Sanctuary/Reserve | 251,050 | 2,761,550 | 7,180,030 | 25 |
| 51 | Botswana | GH/10 | GCA/ WMA / GMA/ CHA | 250,080 | 2,750,880 | 7,152,288 | 381 |
| 52 | Zambia | Liuwa Plain | National park | 249,645 | 2,746,095 | 7,139,847 | 372 |
| 53 | Ethiopia | Gambella | National park | 226,705 | 2,493,755 | 6,483,763 | 958 |
| 54 | Zambia | Machiya-Fungulwe | GCA/ WMA / GMA/ CHA | 221,235 | 2,433,585 | 6,327,321 | 62 |
| 55 | South Sudan | Badingilo | National park | 221,070 | 2,431,770 | 6,322,602 | 1,455 |
| 56 | Tanzania | Serengeti | National park | 214,030 | 2,354,330 | 6,121,258 | 1,776 |
| 57 | Mozambique | Nungo | GCA/ WMA / GMA/ CHA | 213,955 | 2,353,505 | 6,119,113 | 151 |
| 58 | Mozambique | Coutada Oficial No. 12 | GCA/ WMA / GMA/ CHA | 203,980 | 2,243,780 | 5,833,828 | 187 |
| 59 | South Sudan | Shambe | National park | 202,555 | 2,228,105 | 5,793,073 | 218 |
| 60 | Zimbabwe | Chewore | GCA/ WMA / GMA/ CHA | 193,480 | 2,128,280 | 5,533,528 | 253 |
| 61 | Mozambique | Magoe | National park | 192,165 | 2,113,815 | 5,495,919 | 257 |
| 62 | Zambia | Rufunsa | GCA/ WMA / GMA/ CHA | 191,345 | 2,104,795 | 5,472,467 | 219 |
| 63 | Mozambique | Coutada Oficial No. 13 | GCA/ WMA / GMA/ CHA | 188,910 | 2,078,010 | 5,402,826 | 274 |
| 64 | Zambia | Nsumbu | National park | 186,835 | 2,055,185 | 5,343,481 | 85 |
| 65 | Mozambique | Marromeu | Game reserve / National reserve / National sanctuary | 181,530 | 1,996,830 | 5,191,758 | 143 |
| 66 | Botswana | NG/13 | GCA/ WMA / GMA/ CHA | 178,055 | 1,958,605 | 5,092,373 | 143 |
| 67 | Botswana | NG/5 | GCA/ WMA / GMA/ CHA | 170,920 | 1,880,120 | 4,888,312 | 376 |
| 68 | Zimbabwe | Hwange | National park | 168,295 | 1,851,245 | 4,813,237 | 779 |
| 69 | Zambia | Chiawa | GCA/ WMA / GMA/ CHA | 164,350 | 1,807,850 | 4,700,410 | 126 |
| 70 | Mozambique | Coutada Oficial No. 5 | GCA/ WMA / GMA/ CHA | 160,485 | 1,765,335 | 4,589,871 | 416 |
| 71 | Mozambique | Banhine | National park | 153,600 | 1,689,600 | 4,392,960 | 276 |
| 72 | Malawi | Kasungu National Park | National park | 151,845 | 1,670,295 | 4,342,767 | 132 |
| 73 | Tanzania | Mahale Mountains | National park | 149,055 | 1,639,605 | 4,262,973 | 237 |
| 74 | Botswana | GH/11 & GH/13 | GCA/ WMA / GMA/ CHA | 137,300 | 1,510,300 | 3,926,780 | 295 |
| 75 | Botswana | KD/1 | GCA/ WMA / GMA/ CHA | 125,575 | 1,381,325 | 3,591,445 | 262 |
| 76 | Zambia | Lumimba | GCA/ WMA / GMA/ CHA | 114,860 | 1,263,460 | 3,284,996 | 469 |
| 77 | Zimbabwe | Matetsi | GCA/ WMA / GMA/ CHA | 113,840 | 1,252,240 | 3,255,824 | 235 |
| 78 | Zimbabwe | Gonarezhou | National park | 112,670 | 1,239,370 | 3,222,362 | 292 |
| 79 | Zimbabwe | Matusadona | National park | 109,795 | 1,207,745 | 3,140,137 | 104 |
| 80 | Mozambique | Messalo | GCA/ WMA / GMA/ CHA | 104,600 | 1,150,600 | 2,991,560 | 62 |
| 81 | Zimbabwe | Chizarira | National park | 95,805 | 1,053,855 | 2,740,023 | 138 |
| 82 | Zimbabwe | Doma | GCA/ WMA / GMA/ CHA | 92,665 | 1,019,315 | 2,650,219 | 83 |
| 83 | Mozambique | Coutada Oficial No. 11 | GCA/ WMA / GMA/ CHA | 92,625 | 1,018,875 | 2,649,075 | 120 |
| 84 | Tanzania | Udzungwa Mountains | National park | 85,170 | 936,870 | 2,435,862 | 362 |
| 85 | Tanzania | Rungwa | Game reserve / National reserve / National sanctuary | 84,850 | 933,350 | 2,426,710 | 529 |
| 86 | Zimbabwe | Charara | GCA/ WMA / GMA/ CHA | 84,000 | 924,000 | 2,402,400 | 136 |
| 87 | Ethiopia | Chebera Churchura | National park | 81,615 | 897,765 | 2,334,189 | 140 |
| 88 | Zimbabwe | Mana Pools | National park | 79,495 | 874,445 | 2,273,557 | 154 |
| 89 | South Sudan | Meshra | Game reserve / National reserve / National sanctuary | 79,130 | 870,430 | 2,263,118 | 869 |
| 90 | Mozambique | Zinave | National park | 78,825 | 867,075 | 2,254,395 | 232 |
| 91 | Botswana | KW/2 | GCA/ WMA / GMA/ CHA | 78,125 | 859,375 | 2,234,375 | 141 |
| 92 | Tanzania | Kizigo | Game reserve / National reserve / National sanctuary | 77,525 | 852,775 | 2,217,215 | 265 |
| 93 | Zimbabwe | Hurungwe | GCA/ WMA / GMA/ CHA | 73,705 | 810,755 | 2,107,963 | 220 |
| 94 | Botswana | NG/14 | GCA/ WMA / GMA/ CHA | 70,410 | 774,510 | 2,013,726 | 118 |
| 95 | Botswana | NG/4 | GCA/ WMA / GMA/ CHA | 68,665 | 755,315 | 1,963,819 | 100 |
| 96 | Zambia | Chisomo | GCA/ WMA / GMA/ CHA | 68,040 | 748,440 | 1,945,944 | 76 |
| 97 | Botswana | CT/2 | GCA/ WMA / GMA/ CHA | 67,570 | 743,270 | 1,932,502 | 185 |
| 98 | Mozambique | Coutada Oficial No. 4 | GCA/ WMA / GMA/ CHA | 65,310 | 718,410 | 1,867,866 | 209 |
| 99 | Botswana | KD/12 | GCA/ WMA / GMA/ CHA | 64,330 | 707,630 | 1,839,838 | 248 |
| 100 | Democratic Republic of the Congo | Kabobo | Game reserve / National reserve / National sanctuary | 56,010 | 616,110 | 1,601,886 | 136 |
| 101 | Chad | Zakouma | National park | 54,225 | 596,475 | 1,550,835 | 270 |
| 102 | Tanzania | Liparamba | Game reserve / National reserve / National sanctuary | 51,315 | 564,465 | 1,467,609 | 14 |
| 103 | Botswana | Gemsbok | National park | 50,265 | 552,915 | 1,437,579 | 485 |
| 104 | Malawi | Majete Wildlife Reserve | Wildlife Sanctuary/Reserve | 48,870 | 537,570 | 1,397,682 | 71 |
| 105 | Botswana | Makgadikgadi Pans | National park | 48,500 | 533,500 | 1,387,100 | 170 |
| 106 | Botswana | CH/12 | GCA/ WMA / GMA/ CHA | 48,400 | 532,400 | 1,384,240 | 83 |
| 107 | Zambia | Namwala | GCA/ WMA / GMA/ CHA | 48,215 | 530,365 | 1,378,949 | 250 |
| 108 | Uganda | Karuma | Wildlife Sanctuary/Reserve | 47,890 | 526,790 | 1,369,654 | 31 |
| 109 | Botswana | Khutse | Game reserve / National reserve / National sanctuary | 47,670 | 524,370 | 1,363,362 | 75 |
| 110 | Botswana | KD/15 | GCA/ WMA / GMA/ CHA | 42,675 | 469,425 | 1,220,505 | 150 |
| 111 | Zambia | Tondwa | GCA/ WMA / GMA/ CHA | 40,975 | 450,725 | 1,171,885 | 37 |
| 112 | Zimbabwe | Chirisa | GCA/ WMA / GMA/ CHA | 40,910 | 450,010 | 1,170,026 | 116 |
| 113 | Namibia | Mudumu | National park | 38,670 | 425,370 | 1,105,962 | 46 |
| 114 | Tanzania | Ngorongoro | GCA/ WMA / GMA/ CHA | 36,580 | 402,380 | 1,046,188 | 626 |
| 115 | Benin | Pendjari | National park | 36,115 | 397,265 | 1,032,889 | 455 |
| 116 | Botswana | GH/3 | GCA/ WMA / GMA/ CHA | 33,135 | 364,485 | 947,661 | 87 |
| 117 | Zimbabwe | Dande | GCA/ WMA / GMA/ CHA | 31,530 | 346,830 | 901,758 | 43 |
| 118 | Democratic Republic of the Congo | Luama-Katanga | GCA/ WMA / GMA/ CHA | 31,040 | 341,440 | 887,744 | 88 |
| 119 | Botswana | NG/18 | GCA/ WMA / GMA/ CHA | 29,920 | 329,120 | 855,712 | 72 |
| 120 | Ethiopia | Bale Mountains | National park | 29,775 | 327,525 | 851,565 | 92 |
| 121 | Botswana | GH/2 | GCA/ WMA / GMA/ CHA | 29,695 | 326,645 | 849,277 | 109 |
| 122 | United Republic of Tanzania | Saadani | National park | 28,110 | 309,210 | 803,946 | 54 |
| 123 | Botswana | SO/2 | GCA/ WMA / GMA/ CHA | 27,595 | 303,545 | 789,217 | 72 |
| 124 | South Africa | Marakele National Park | National park | 27,575 | 303,325 | 788,645 | 36 |
| 125 | Zimbabwe | Sapi | GCA/ WMA / GMA/ CHA | 27,430 | 301,730 | 784,498 | 83 |
| 126 | South Africa | Hluhluwe-Imfolozi Game Reserve | Game reserve / National reserve / National sanctuary | 26,950 | 296,450 | 770,770 | 124 |
| 127 | Tanzania | Tarangire | National park | 26,520 | 291,720 | 758,472 | 182 |
| 128 | Tanzania | Burigi | Game reserve / National reserve / National sanctuary | 26,085 | 286,935 | 746,031 | 232 |
| 129 | Botswana | Nxai Pan | National park | 25,895 | 284,845 | 740,597 | 113 |
| 130 | Malawi | Liwonde National Park | National park | 25,050 | 275,550 | 716,430 | 64 |
| 131 | Botswana | NG/16 | GCA/ WMA / GMA/ CHA | 24,580 | 270,380 | 702,988 | 63 |
| 132 | Botswana | CT/1 | GCA/ WMA / GMA/ CHA | 23,870 | 262,570 | 682,682 | 161 |
| 133 | Uganda | Toro-Semuliki or Toro-Semliki | Wildlife Sanctuary/Reserve | 22,620 | 248,820 | 646,932 | 49 |
| 134 | Malawi | Vwaza Marsh Wildlife Reserve | Wildlife Sanctuary/Reserve | 22,480 | 247,280 | 642,928 | 48 |
| 135 | Botswana | Moremi | Game reserve / National reserve / National sanctuary | 22,390 | 246,290 | 640,354 | 223 |
| 136 | Mozambique | Chimanimani | Wildlife Sanctuary/Reserve | 22,140 | 243,540 | 633,204 | 34 |
| 137 | Mozambique | Maputo | Wildlife Sanctuary/Reserve | 21,905 | 240,955 | 626,483 | 82 |
| 138 | Botswana | KD/2 | GCA/ WMA / GMA/ CHA | 21,835 | 240,185 | 624,481 | 163 |
| 139 | Botswana | NG/41 | GCA/ WMA / GMA/ CHA | 21,770 | 239,470 | 622,622 | 105 |
| 140 | Botswana | KW/4 | GCA/ WMA / GMA/ CHA | 21,620 | 237,820 | 618,332 | 31 |
| 141 | Zambia | Sichifula | GCA/ WMA / GMA/ CHA | 21,280 | 234,080 | 608,608 | 177 |
| 142 | Botswana | NG/20 | GCA/ WMA / GMA/ CHA | 17,795 | 195,745 | 508,937 | 83 |
| 143 | Kenya | Masai Mara | Game reserve / National reserve / National sanctuary | 17,515 | 192,665 | 500,929 | 263 |
| 144 | Botswana | NG/49 | GCA/ WMA / GMA/ CHA | 16,435 | 180,785 | 470,041 | 42 |
| 145 | Botswana | NG/42 | GCA/ WMA / GMA/ CHA | 15,640 | 172,040 | 447,304 | 138 |
| 146 | Kenya | Tsavo East and West | National park | 14,525 | 159,775 | 415,415 | 1,169 |
| 147 | Botswana | NG/43 | GCA/ WMA / GMA/ CHA | 13,670 | 150,370 | 390,962 | 157 |
| 148 | Zimbabwe | Zambezi | National park | 13,295 | 146,245 | 380,237 | 34 |
| 149 | Kenya | Chyulu Hills | National park | 12,150 | 133,650 | 347,490 | 35 |
| 150 | Botswana | CT/3 | GCA/ WMA / GMA/ CHA | 11,070 | 121,770 | 316,602 | 84 |
| 151 | Ethiopia | Mago | National park | 10,425 | 114,675 | 298,155 | 192 |
| 152 | Botswana | NG/47 | GCA/ WMA / GMA/ CHA | 10,290 | 113,190 | 294,294 | 76 |
| 153 | Zambia | Sandwe | GCA/ WMA / GMA/ CHA | 9,165 | 100,815 | 262,119 | 133 |
| 154 | Botswana | NG/34 | GCA/ WMA / GMA/ CHA | 9,030 | 99,330 | 258,258 | 26 |
| 155 | Tanzania | Swaga Swaga | Game reserve / National reserve / National sanctuary | 8,975 | 98,725 | 256,685 | 54 |
| 156 | Botswana | KD/6 | GCA/ WMA / GMA/ CHA | 8,280 | 91,080 | 236,808 | 43 |
| 157 | South Africa | Kalahari Gemsbok National Park | National park | 7,655 | 84,205 | 218,933 | 133 |
| 158 | Cameroon | Waza | National park | 6,215 | 68,365 | 177,749 | 107 |
| 159 | Namibia | Nkasa Rupara | National park | 5,435 | 59,785 | 155,441 | 14 |
| 160 | Kenya | Dodori | Game reserve / National reserve / National sanctuary | 5,010 | 55,110 | 143,286 | 31 |
| 161 | Tanzania | Muhesi | Game reserve / National reserve / National sanctuary | 4,885 | 53,735 | 139,711 | 221 |
| 162 | Botswana | KD/5 | GCA/ WMA / GMA/ CHA | 4,785 | 52,635 | 136,851 | 16 |
| 163 | Botswana | NG/15 | GCA/ WMA / GMA/ CHA | 4,680 | 51,480 | 133,848 | 50 |
| 164 | Botswana | KW/12 | GCA/ WMA / GMA/ CHA | 4,385 | 48,235 | 125,411 | 6 |
| 165 | Kenya | Boni | Game reserve / National reserve / National sanctua | 4,210 | 46,310 | 120,406 | 52 |
| 166 | Botswana | CT/10 | GCA/ WMA / GMA/ CHA | 4,205 | 46,255 | 120,263 | 32 |
| 167 | Botswana | KW/6 | GCA/ WMA / GMA/ CHA | 3,395 | 37,345 | 97,097 | 15 |
| 168 | United Republic of Tanzania | Mkungunero | Game reserve / National reserve / National sanctuary | 3,010 | 33,110 | 86,086 | 36 |
| 169 | Burkina Faso | Arli | National park | 2,960 | 32,560 | 84,656 | 890 |
| 170 | Botswana | CH/11 | GCA/ WMA / GMA/ CHA | 2,925 | 32,175 | 83,655 | 50 |
| 171 | Botswana | NG/19 | GCA/ WMA / GMA/ CHA | 2,875 | 31,625 | 82,225 | 8 |
| 172 | Botswana | NG/24 | GCA/ WMA / GMA/ CHA | 2,720 | 29,920 | 77,792 | 28 |
| 173 | Kenya | Meru | National park | 2,450 | 26,950 | 70,070 | 34 |
| 174 | Botswana | CT/11 | GCA/ WMA / GMA/ CHA | 2,090 | 22,990 | 59,774 | 96 |
| 175 | Tanzania | Kilimanjaro | National park | 2,055 | 22,605 | 58,773 | 251 |
| 176 | South Africa | Pilanesberg National Park | ZAF Nature reserve | 2,000 | 22,000 | 57,200 | 66 |
| 177 | South Africa | Addo-Elephant National Park | National park | 1,425 | 15,675 | 40,755 | 48 |
| 178 | Botswana | KD/11 | GCA/ WMA / GMA/ CHA | 1,245 | 13,695 | 35,607 | 8 |
| 179 | Tanzania | Mkomazi | National park | 1,180 | 12,980 | 33,748 | 187 |
| 180 | Uganda | Lake Mburo | National park | 945 | 10,395 | 27,027 | 16 |
| 181 | Zimbabwe | Malipati | GCA/ WMA / GMA/ CHA | 850 | 9,350 | 24,310 | 6 |
| 182 | Namibia | Hobatere | Game reserve / National reserve / National sanctua | 790 | 8,690 | 22,594 | 5 |
| 183 | Ethiopia | Maze | National park | 630 | 6,930 | 18,018 | 51 |
| 184 | Botswana | NG/21 | GCA/ WMA / GMA/ CHA | 585 | 6,435 | 16,731 | 8 |
| 185 | Kenya | North Kitui | Game reserve / National reserve / National sanctuary | 550 | 6,050 | 15,730 | 30 |
| 186 | Zimbabwe | Chete | GCA/ WMA / GMA/ CHA | 515 | 5,665 | 14,729 | 93 |
| 187 | Ethiopia | Babille | Wildlife Sanctuary/Reserve | 500 | 5,500 | 14,300 | 368 |
| 188 | Ethiopia | Nechisar | National park | 390 | 4,290 | 11,154 | 21 |
| 189 | Kenya | Marsabit National Park & Reserve | National park | 340 | 3,740 | 9,724 | 90 |
| 190 | Ethiopia | Yabello | Wildlife Sanctuary/Reserve | 330 | 3,630 | 9,438 | 144 |
| 191 | Ethiopia | Mille-Sardo | Wildlife Sanctuary/Reserve | 250 | 2,750 | 7,150 | 148 |
| 192 | Kenya | Arawale | Game reserve / National reserve / National sanctua | 215 | 2,365 | 6,149 | 28 |
| 193 | Ethiopia | Kafa | Biosphere Reserve | 190 | 2,090 | 5,434 | 23 |
| 194 | Ethiopia | Geralle | National park | 125 | 1,375 | 3,575 | 153 |
| 195 | Tanzania | Biharamulo | Game reserve / National reserve / National sanctua | 120 | 1,320 | 3,432 | 35 |
| 196 | Kenya | Shaba | Game reserve / National reserve / National sanctuary | 90 | 990 | 2,574 | 12 |
| 197 | Kenya | Amboseli | National park | 5 | 55 | 143 | 18 |
| 198 | Kenya | Samburu | Game reserve / National reserve / National sanctuary | 0 | 0 | 0 | 6 |
|  | Total |  |  | 59,628,060 | 655,908,660 | 1,705,362,516 | 62,277 |
|  | Mean |  |  | 301,152 | 3,312,670 | 8,612,942 | 315 |
|  | SD |  |  | 875,162 | 9,626,784 | 25,029,639 | 544 |
|  | N |  |  | 198 | 198 | 198 | 198 |

**Table S3**. Highest combined priority ranking of lion protected areas (PA) for both potential carbon revenue (PCR) generation and predicted lion population size (i.e., carrying capacity).

| Combined Rank | Combined score | Africa Lion PA | Country | Predicted Lion Population Size | Low PCR Estimate (USD $/yr) | High PCR Estimate (USD $/yr) |
| --- | --- | --- | --- | --- | --- | --- |
| 1 | 2 | Luengue-Luiana | Angola | 5,633 | 7,788,312 | 22,745,714 |
| 3 | 5 | Niassa | Mozambique | 1,881 | 6,765,565 | 193,495,153 |
| 3 | 5 | Selous | Tanzania | 2,157 | 4,153,721 | 118,796,419 |
| 4 | 12 | Kafue | Zambia | 1,714 | 1,564,726 | 44,751,166 |
| 5 | 13 | Manovo-Gounda-Saint Floris | Central Africa Republic | 1,457 | 3,993,089 | 114,202,353 |
| 6 | 18 | Musalangu | Zambia | 1,594 | 1,242,875 | 35,546,217 |
| 7 | 20 | Central Kalahari | Botswana | 1,616 | 918,503 | 26,269,243 |
| 8 | 22 | South Luangwa | Zambia | 1,104 | 1,260,615 | 36,053,589 |
| 9 | 30 | West Zambezi | Zambia | 1,254 | 642,915 | 18,387,369 |
| 10 | 36 | Luano | Zambia | 800 | 702,844 | 20,101,341 |
| Median |  |  |  | 1,605 | 1,412,671 | 35,799,903 |
| Total |  |  |  | 19,210 | 29,033,165 | 630,348,564 |

| **a** 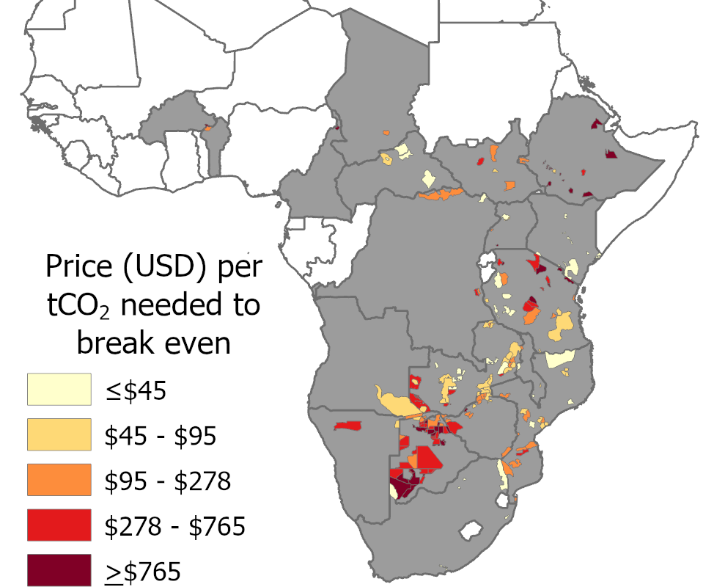 | **b** 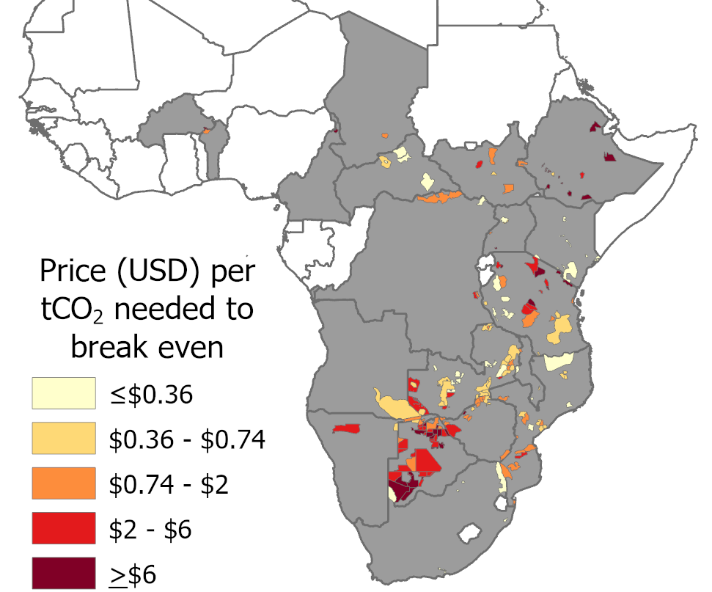 |
| --- | --- |

**Figure S1.** The market price needed to cover the estimated funding gap in each of 198 protected areas (based on Lindsey et al.^12^) with >0 GHG abatement potential if the revenue were generated by carbon projects. The estimates show the difference between selling carbon credits from a) emissions reduction alone (e.g., Lipsett-Moore et al.^31^), and b) multiple methods that combine possible credits from four existing fire-driven carbon methodologies that would add carbon sequestration with emissions reductions. Nineteen countries (gray) contain at least 1 of the 198 PAs.

|  | **a** Early Dry Season  Average Emissions | **b** Late Dry Season  Average Emissions |
| --- | --- | --- |
| Woody Biomass  (mg/ha) | 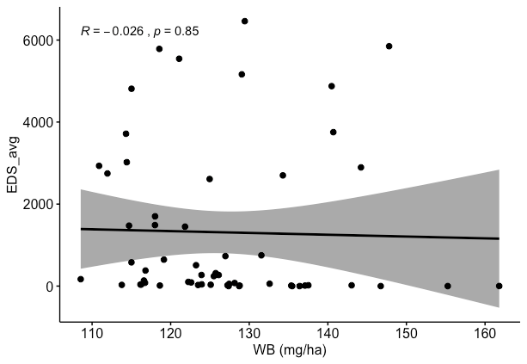 | 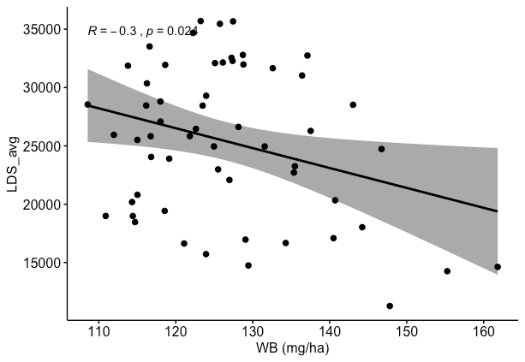 |
| Fire Frequency  (mean number of fires) | 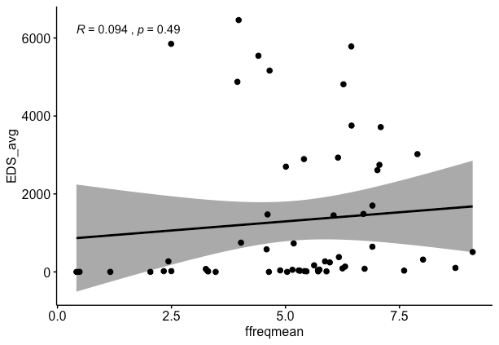 | 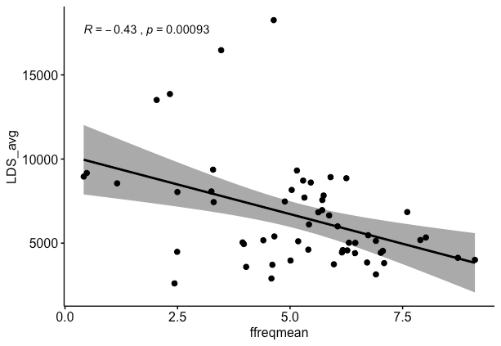 |
|  | **c** Early Dry Season Emissions Variability ( (Std. Dev.) | **d** Late Dry Season Emissions Variability (Std. Dev.) |
| Woody Biomass  (mg/ha) | 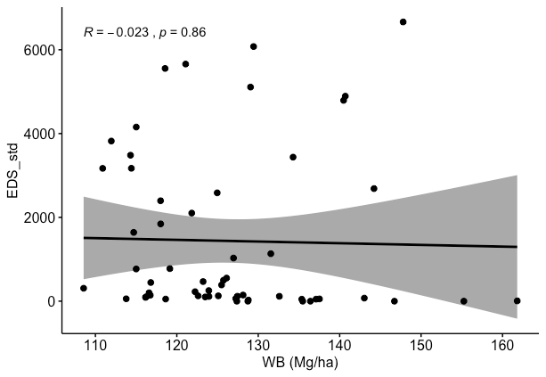 | 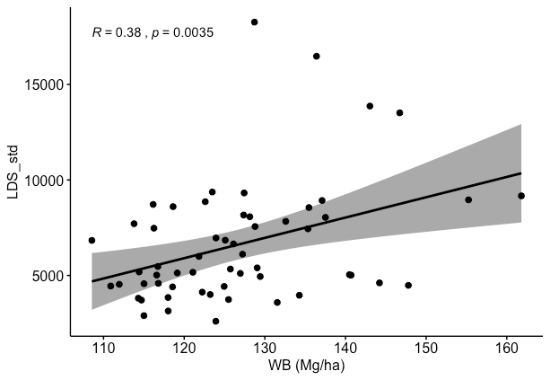 |
| Fire Frequency  (mean number of fires) | 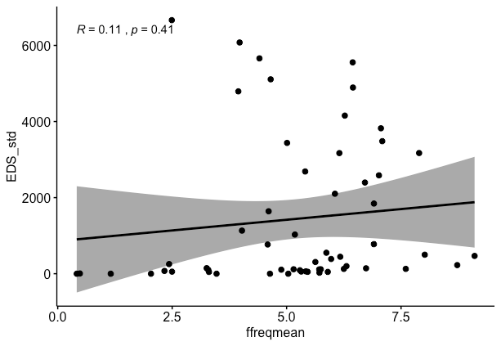 | 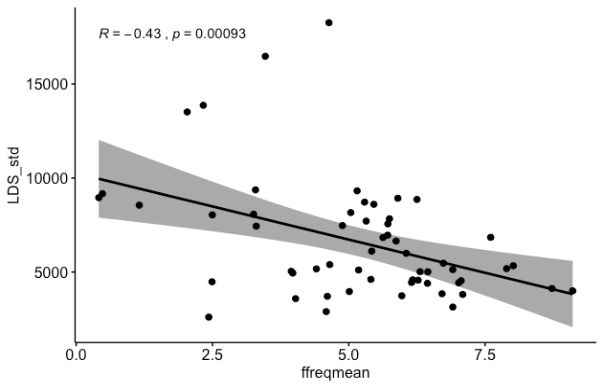 |

**Figure S2.** Linear regression relationships in Niassa National Reserve on a per-pixel basis for a) early (EDS) emissions, b) late (LDS) emissions, c) EDS variability and d) LDS variability in relation to mean fire frequency (number of fires) and mean woody biomass (mg/ha).

In Niassa, higher average emissions were associated with greater mean number of fires (p<0.001) and less woody biomass (mg/ha) (p<0.02), suggesting LDS fires burned hotter and covered more area to produce greater emissions. In contrast, there were no significant relationships (p>0.05) among emissions or their variability for either fire frequency or woody biomass with EDS fires, suggesting that early fires were “cool” and/or patchy enough that they had weaker, if any effects on woody carbon and emissions. This supports the prediction that LDS fires in Niassa would create more emissions and be more damaging than EDS fires. A fire management program designed to reduce this risk would generate similar PCR benefits predicted by the Australian EDS approach.
